# Integrated multi-omics profiling uncovers the epigenetic, transcriptional, and metabolic landscape of prostate cancer progression

**DOI:** 10.64898/2026.02.11.704810

**Authors:** Christine Aaserød Pedersen, Maximilian Wess, Maria K. Andersen, Elise Midtbust, Abhibhav Sharma, Thomas Fleischer, Morten Beck Rye, May-Britt Tessem

## Abstract

A comprehensive understanding of the underlying molecular mechanisms of prostate cancer is essential for the development of precise diagnostic biomarkers. In this study, we applied unsupervised multi-omics factor analysis (MOFA) to integrate DNA methylation, gene expression, and metabolic profiles derived from the same individuals, aiming to characterize the biological landscape of normal, malignant, and aggressive prostate tissue. Our analysis identified distinct molecular pathways associated with aggressive disease, specifically those involved in zinc metabolism, cell cycle regulation, smooth muscle architecture, immune activation, and tissue morphology. Key metabolites within the TCA cycle, amino acid metabolism, and lipid pathways were central to these signatures. Furthermore, we observed a consistent co-enrichment of SP1 and CTCFL binding regions among factor-associated CpGs, suggesting a model of global epigenetic reprogramming. These findings indicate a novel interplay between Polycomb deregulation, CTCFL-mediated chromatin remodeling, and SP1-driven transcriptional activation in shaping the prostate cancer epigenome. Apart from immune activation, the identified molecular signatures were validated in the TCGA cohort and demonstrated significant predictive value for disease recurrence. Overall, these results underscore the power of multi-omics integration in providing a holistic understanding of prostate cancer biology and its potential for clinical translation into prognostic biomarkers.

## Introduction

Prostate cancer is one of the most prevalent cancers globally, affecting approximately one in eight men (1, 2). The current diagnostic pathway struggles to accurately differentiate between patients with indolent and aggressive disease. This clinical challenge necessitates improved stratification methods to minimize unnecessary treatment side effects for indolent cases and ensure early intervention for aggressive cases.

Understanding which pathways that lead to aggressive prostate cancer is needed for identifying biomarkers and drug targets. However, newer tissue-based molecular biomarker tools have shown inconsistent performance due to molecular and tissue heterogeneity (3). Multi-omics analysis, which examines multiple omics layers simultaneously from the same samples, may elucidate complex mechanisms and molecular interactions, providing a deeper understanding of disease progression. Studies have shown that combining features from multiple omics layers enhances biomarker precision compared to single-omics approaches (4, 5).

This study utilized multi-omics factor analysis (MOFA), a versatile unsupervised machine learning tool that integrates various omics datasets into factors representing patterns of biological variation (6). We included integrated metabolomics, transcriptomics and DNA methylation data into the MOFA model. Metabolites are integral to biochemical pathways and closely linked to cell phenotype, while transcriptomics data reflect gene activity and regulation. Lastly, the DNA methylation data covers important epigenetic modification crucial for regulating gene expression. Together, these three omics levels can shed a unique light on disease development at the cellular level.

The role of DNA methylation in gene expression and cell differentiation is complex. DNA methylation regulates chromatin structure by inhibiting or recruiting protein complexes that affect chromatin packing. One such complex is the polycomb repressive complex (PRC), which is implicated in prostate cancer aggressiveness and regulates cell differentiation through histone modifications, impacting chromatin accessibility for transcription (7, 8). DNA methylation also influences transcription factor affinity to DNA and the transcriptional machinery’s affinity to genes, of which the most common and most studied is promoter methylation. Cell metabolism is also closely linked to methylation by i.e. regulating the availability of methyl groups required for methylation (9). Given the complexity and tissue-specific nature of these processes, studying various aspects of DNA methylation alongside gene expression and metabolites is essential for understanding these intricate biological mechanisms.

Prostate cancer research faces major challenges due to combined tissue and molecular heterogeneity, making it difficult to pinpoint the molecular mechanisms involved in aggressive disease. Comprehensive research using tissues from different prostate locations and integrating information across molecular layers is required to understand these mechanisms. Our study aims to capture the molecular complexity of tissue heterogeneity across omics levels and assess the predictive value of integrated multi-omics analysis of methylation, gene expression, and metabolites for cancer aggressiveness.

## Materials and methods

We included in total 280 samples from 57 patients. All patients underwent radical prostatectomy (RP) as curative treatment for prostate cancer at St Olavs Hospital, Trondheim, Norway, between 2007 and 2017. Immediately after RP, a 2 mm thick slice from the middle of the prostate was collected, frozen and stored at -80 °C by Biobank1, the research biobank of mid-Norway following the protocol established by Bertilsson *et al*. (10). All patients delivered a written informed consent, and the study was approved by the Norwegian regional ethical committee (2017/576), which follows EU and Norwegian ethical regulations.

Three to five samples of 3mm in diameter were collected from each frozen prostate slice, aiming to sample both tumor, normal and normal adjacent tissue, as shown in Figure 1. After collection, the tissue type (normal or cancer) and eventual cancer grade was annotated by a histopathologist using a hematoxylin, erythrosine, saffron (HES)-stained section from the sample. Stroma and cancer content were estimated based on histopathology. Clinical follow-up data was collected 6-16 years after radical prostatectomy. Biochemical recurrence was defined as prostate-specific antigen (PSA) above 0.2ng/ml after radical prostatectomy. In the “recurrence” patient group (N = 45), we also included patients that had persistent PSA after surgery (PSA>0.1ng/ml), meaning the PSA remained detectable. Patients in the “metastasis” group (N = 29) were diagnosed with metastatic disease by clinical imaging during follow-up, and patients in the “castrate resistant prostate cancer” group (N = 13) had elevated PSA levels after receiving androgen deprivation treatment after radical prostatectomy. The median follow-up time was 9.2 years (IQR 8.25-12.3 years). Sample characteristics are described in Table 1.

**Table 1.** Sample characteristics for methylation, transcriptomics and metabolomics. IQR: interquartile range.

|  | Methylation |  | Transcriptomics |  | Metabolites |  | Total |
| --- | --- | --- | --- | --- | --- | --- | --- |
|  | Cancer (N=51) | Normal (N=45) | Cancer (N=114) | Normal (N=62) | Cancer (N=184) | Normal (N=94) | Total (N=280) |
| **Sample type, n (%)** |  |  |  |  |  |  |  |
| - ISUP1 | 13 (25.5%) | - | 32 (28.1%) | - | 61 (33.2%) | - | 61 (21.8%) |
| - ISUP2 | 19 (37.3%) | - | 30 (26.3%) | - | 48 (26.1%) | - | 48 (17.1%) |
| - ISUP3 | 10 (19.6%) | - | 21 (18.4%) | - | 29 (15.8%) | - | 31 (11.1%) |
| - ISUP4 | 5 (9.8%) | - | 20 (17.5%) | - | 32 (17.4%) | - | 32 (11.4%) |
| - ISUP5 | 4 (7.8%) | - | 11 (9.6%) | - | 14 (7.6%) | - | 14 (5.0%) |
| - Normal | - | 23 (51.1%) | - | 35 (56.5%) | - | 51 (54.3%) | 51 (18.2%) |
| - Normal adjacent | - | 22 (48.9%) | - | 27 (43.5%) | - | 43 (45.7%) | 43 (15.4%) |
| **Cancer content** |  |  |  |  |  |  |  |
| - Median | 93.0 | - | 76.5 | - | 71.0 | - | 30.0 |
| - IQR | 70.5 - 100 | - | 49.2 - 100 | - | 30 - 100 | - | 0.0, 90.2 |
| **Stroma content** |  |  |  |  |  |  |  |
| - Median | 25.0 | 60.0 | 35.0 | 60.0 | 35.0 | 62.5 | 47.5 |
| - IQR | 17.5 - 35 | 50 - 70 | 20 - 50 | 50 - 65 | 20 - 50 | 50 - 70 | 30 - 65 |
| **EAU risk group, n (%)** |  |  |  |  |  |  |  |
| - High | 17 (33.3%) | 15 (33.3%) | 51 (44.7%) | 25 (40.3%) | 91 (49.5%) | 43 (45.7%) | 134 (47.9%) |
| - Intermediate | 34 (66.7%) | 30 (66.7%) | 63 (55.3%) | 37 (59.7%) | 93 (50.5%) | 51 (54.3%) | 146 (52.1%) |
| **Recurrence, n (%)** |  |  |  |  |  |  |  |
| - Non-recurrence | 18 (35.3%) | 14 (31.1%) | 24 (21.1%) | 16 (25.8%) | 35 (19.0%) | 22 (23.4%) | 58 (20.7%) |
| - Recurrence | 33 (64.7%) | 31 (68.9%) | 90 (78.9%) | 46 (74.2%) | 149 (81.0%) | 72 (76.6%) | 222 (79.3%) |
| **Metastasis, n (%)** |  |  |  |  |  |  |  |
| - No | 35 (68.6%) | 30 (66.7%) | 61 (53.5%) | 35 (56.5%) | 95 (51.6%) | 51 (54.3%) | 147 (52.5%) |
| - Yes | 16 (31.4%) | 15 (33.3%) | 53 (46.5%) | 27 (43.5%) | 89 (48.4%) | 43 (45.7%) | 133 (47.5%) |
| **Castrate resistance, n (%)** |  |  |  |  |  |  |  |
| - No | 45 (88.2%) | 40 (88.9%) | 75 (67.6%) | 47 (77.0%) | 132 (73.7%) | 74 (79.6%) | 207 (75.5%) |
| - Yes | 6 (11.8%) | 5 (11.1%) | 36 (32.4%) | 14 (23.0%) | 47 (26.3%) | 19 (20.4%) | 67 (24.5%) |

**Figure 1:**
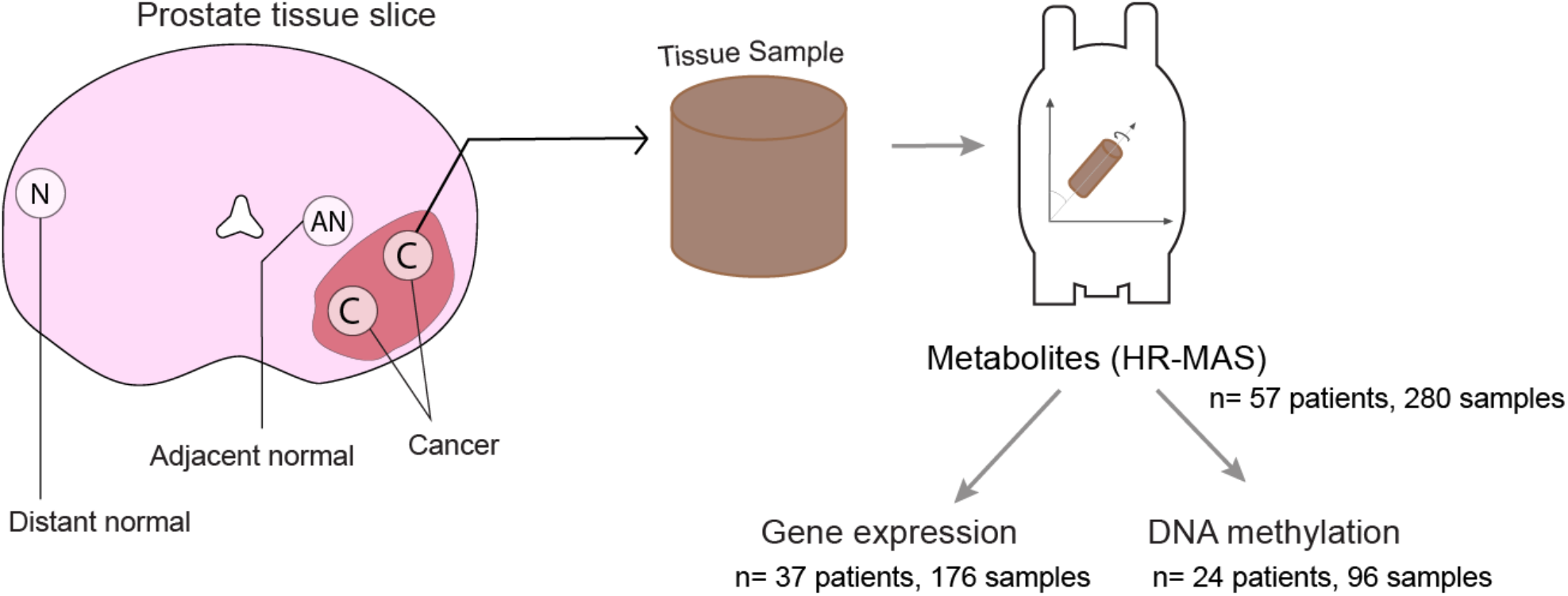
Sample collection and analysis. From each patient, 3-5 samples were collected. One sample was extracted far away from the tumor, one normal sample adjacent to the tumor, and two cancer samples were collected from the tumor area. The samples were drilled from the fresh frozen tissue slice and resembled cylinders in shape. The samples were analyzed with HRMAS (High-Resolution Magic Angle Spinning) for metabolite detection, and then DNA and RNA was extracted and analyzed for gene expression and DNA methylation, respectively.

### HR-MAS NMR/metabolite data

All samples were first analyzed by HR-MAS (high resolution - magic angle spinning), an NMR spectroscopy method leaving the tissue intact for further analysis. It was performed on a 600 MHz Bruker Avance DRX600 (14.1 T) III NMR spectrometer (Bruker BioSpin, Germany) using the pulse sequence *noesygppr1d* on a ^1^H/^13^C MAS probe.

The raw spectra were preprocessed using python version 3.8. After Automics phasing (11), the water peak visible from to 4.6ppm to 6ppm was removed, spectra were baseline corrected using asymmetric least squares smoothing (12) and aligned using icoshift (13). Further, the peaks were mean matrix normalized. Peaks were picked and identified manually based on previous knowledge of HRMAS metabolites in prostate tissue (14-16), resulting in 30 identified metabolites. We performed relative quantification by integrating the area of the selected peaks. In case of several peaks per metabolite, we used the one that had the least overlap with other compound peaks. Two samples were removed during preprocessing due to noisy spectra because of high lipid levels, leaving 278 samples for analysis.

### Transcriptomics data

DNA and RNA was extracted from the tissue after HRMAS analysis, using the QIAGEN AllPrep DNA/RNA/miRNA kit. RNA from 176 samples was sequenced using the cDNA library SENSE mRNA-Seq Library Prep Kit V2 (600 ng RNA as input). Single-read sequencing was performed using the NextSeq 500/550 High Output kit v2.5 (75 cycles) on an Illumina NextSeq 500 instrument.

Produced FASTQ files were filtered and trimmed using fastp v0.20.0. The sequence alignment was performed using the STAR tool against the reference genome set GRCh38 release 92 (Ensembl). Subsequently, *featureCounts* (17) was used to extract gene counts from sequence reads according to the reference set.

RNA was sequenced in 4 different batches on different dates, and a batch effect was identified in exploratory analyses. The batch effect was adjusted for using ComBat-seq (18). Further, the data was normalized for library size, and CPM (counts per million) values were extracted and log transformed.

### DNA methylation preprocessing

The isolated DNA from 96 samples was bisulfite converted using the EZ DNAMethylation kit, and the converted DNA was analyzed for methylation using the Illumina HumanMethylation EPIC BeadChip. For 64 of the samples, we used the Illumina EPIC array, and for 32 samples we used the EPIC v2 array.

We filtered out the probes with a detection p value <0.05 and a bead count of under 3 in more than 5% of samples. Probes not measuring a CpG were removed, as were probes where the CpGs were closer less than 5 base pairs away from known SNPs, to avoid potential confounding by genetic variation (19).

Raw DNA methylation data was preprocessed using the R-package *minfi* and the funnorn-function was used for functional normalization (20). The EPIC v1 and EPICv2 data was combined into one dataset including only the probes with matching probe sequence. A batch effect was observed between samples from the two arrays, that was corrected using combat (21). We used methylation M values as input in the analysis and beta values for plotting, as recommended by Du *et al* (22).

### Multiomics factor analysis

For multi-omics analysis we used the Multi-omics factor analysis (MOFA) tool (6) which is a an unsupervised machine learning model based on factor analysis tailored for handling multi-omics data when the omics levels are analyzed in the same sample. We used the whole metabolite data set (278 samples, 30 metabolites), the top 5000 variable genes and top 5000 variable CpGs from the transcriptomics dataset (176 samples) and methylation dataset (96 samples) based on the package authors recommendation (https://biofam.github.io/MOFA2/faq.html). We trained the MOFA model for 10 factors, since more factors did not increase the explained variance of the model (Supplementary Figure 1). The MOFA model assumes linear mappings from latent factors to observed molecular measurements (6).

### Gene set enrichment

Using the MOFA package built-in function, we performed gene set enrichment analysis (GSEA) for each of the Factors 1-5 once using the positive weighted genes and once using the negative weighted genes. We used the gene sets from msigdb c5 (gene ontology) and extracted the top 5 GO terms with p <0.01 (fdr) for the positive and negative weights for all factors. Further, GO terms were manually grouped together to reflect biological themes.

### Chromatin state assessment

To explore how the CpGs are related to chromatin states in prostate cancer tissue, we used ChromHMM data from the prostate cancer cell lines PC3 and LNCaP. ChromHMM is a software for learning and characterizing chromatin states to functionally annotate the genome by using a multivariate Hidden Markov Model (HMM) for identifying combinatorial patterns of histone marks obtained from ChIP-seq data (23). ChromHMM segmentation data were downloaded from the ENCODE data portal (24) from two prostate cancer cell lines: PC-3 (Accession ENCSR103ZFL) and LNCaP (Accession ENCSR072ZGM). Functional regions were intersected with CpG positions using BEDTools v2.27.1 (25).

Enrichment of CpGs in a ChromHMM defined functional region was measured as the ratio between the frequency of the top (positive) quartile and bottom (negative) quartile of CpGs in each factor over the frequency of CpGs from the Illumina HumanMethylationEPIC array found within the same region. P-values were obtained by hypergeometric testing with the Illumina EPIC array probes as background. Multiple testing was accounted for using Benjamini-Hochberg correction.

### Enrichment for transcription factor binding regions

Maps of direct transcription factor-DNA (TF-DNA) interactions were obtained from the UniBind database (https://unibind.uio.no) (26). The UniBind TFBSs (transcription factor binding sites) represent high-confidence direct TF-DNA interactions determined experimentally through ChIP-seq and computationally through position weight matrices (PWMs) from JASPAR (27). The predicted transcription factor binding sites (TFBSs) were derived from 4114 ChIP-seq experiments for 268 TFs across cell types and tissues and were predicted to have high PWM scores and be near ChIP-seq peaks. TFBSs were extended by ±150bp (then called transcription factor binding regions; TFBRs) and were intersected with CpG positions using BEDTools v2.27 (25). Since UniBind maps of TF binding sites are often derived from several ChIP-seq experiments for each TF, we merged the TF binding sites for all ChIP-seq experiments for each TF. Enrichment of CpGs in TFBRs was imputed using hypergeometric testing (R function phyper) using the IlluminaMethylationEPIC CpGs as background. Multiple testing was accounted for using Benjamini-Hochberg correction.

### Gene signature construction and validation of prognostic impact

We constructed gene signatures for each of 5 biological themes (cell cycle, metallothionein, immune response, tissue morphology and smooth muscle; Figure 2). For a given theme, we selected genes that were involved in the theme’s GO-terms and in the upper or lower quartile of the density distribution of weights in the MOFA factors associated with the theme.

**Figure 2:**
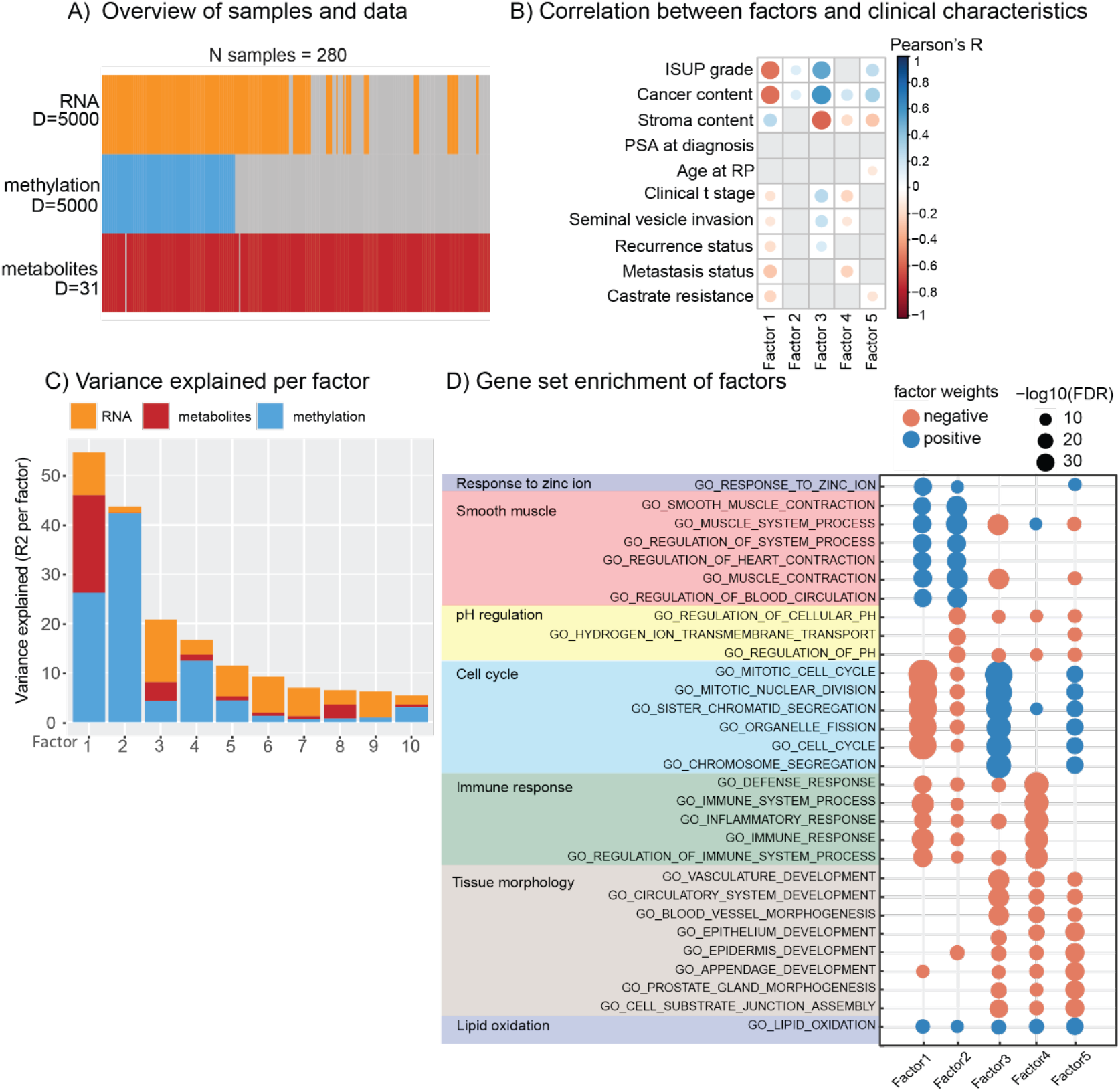
MOFA model reveals biology associated with tissue morphology and clinical outcome. **A)** We included a total of 280 samples, incorporating 5000 genes (RNA), 5000 CpGs (methylation), and 31 metabolites into the model. Gray cells indicate that an omics type was not available for a patient. **B**) Correlation between factors, and clinical and sample characteristics show that the factors are both correlated to sample characteristics such as ISUP grade and stroma content, but also clinical outcomes such as recurrence, metastasis and castrate resistance. **C)** Contribution of each -omic level to the variance explained by the factors generated by MOFA. **D)** Gene set enrichment of the genes in each factor, positive and negative weights analyzed separately. Several biological themes emerged, including response to zinc ion, smooth muscle, pH regulation, cell cycle, immune response, tissue morphology, and lipid oxidation. MOFA = Multi-Omics Factor Analysis; ISUP = International Society for Urological Pathology; PSA = Prostate-specific Antigen; RP = Radical Prostatectomy; FDR = False Discovery Rate

To evaluate the prognostic relevance of gene signatures, we performed single-sample gene set enrichment analysis (ssGSEA; GSVA package)(28) using bulk RNA-seq data from TCGA-PRAD. The bulk RNA-seq expression data were obtained from cbioportal database (https://www.cbioportal.org/datasets) under the heading Prostate Adenocarcinoma (TCGA, PanCancer Atlas). Clinical metadata for this TCGA-PRAD (portal.gdc.cancer.gov/projects/TCGA-PRAD) cohort were curated to construct a harmonized metadata table. The status of biochemical recurrence (BCR) was derived from the NEW_TUMOR_EVENT_AFTER_INITIAL_TREATMENT field, with entries labeled “Yes”, cases annotated as “1:Recurred/Progressed” in DFS_STATUS, or “1:DEAD WITH TUMOR” in DSS_STATUS. The resulting metadata were merged with the bulk TCGA-PRAD RNA-seq data and used for Kaplan-Meier and log-rank survival analyses to evaluate the impact of gene profiles on BCR. Each patient received an enrichment score per signature, which was stratified into low, medium, and high categories (low < 30%, high > 70%, medium 30-70% quantiles). Recurrence-free survival was then analyzed using Kaplan-Meier and log-rank tests across signature-defined groups, visualized using *survminer* in R.(29)

### Feature selection and statistical analysis

To focus on the most dominant patterns in MOFA’s computed factors, we selected genes from GO terms that also had factor weights in the upper or lower quartile of the density distribution of the involved factor’s weights for each biological theme. Some GO terms contain large sets of genes, many of which overlap with other GO terms. For our analyses, we concentrated on genes with functions most relevant to the prostate, so only this selected subset is shown rather than all genes associated with each term. Similarly, we selected metabolites if their associated factor weight was in the upper or lower quantile of the density distribution of the involved factor’s weights. To describe the relationship of the various omics layers, we further selected genes, metabolites, enriched chromatin states, and TFBSs, where genes were significantly correlated (absolute Spearman’s rho > 0.45) with any other omics measurement. Further, we assessed whether expression levels of multi-omics data show clinical significance using Spearman correlation.

## Results and discussion

### Multi-omics factor analysis reveals biological themes associated with clinical outcomes

To reveal the intricate connections between transcriptomics, metabolomics, and epigenomics data, we used MOFA, an unsupervised factor analysis framework for multi-omics data. We used the 5000 most variable genes, 5000 most variable CpGs, and 31 metabolites from 280 samples (N= 57 patients) to train a model with 10 factors (Figure 2A). Each trained factor is associated with the weight for each gene, metabolite, and CpG. A high positive weight reflects a strong association between a variable (i.e., a gene, metabolite, or CpG) and a factor, while a high negative weight reflects a strong inverse relationship. The trained model explained 76% of the total variance in methylation, 50% of the RNA, and 29% of the variance in metabolites (R^2^).

The factors of the MOFA model captured various aspects of prostate cancer-specific clinical and morphological variation (Figure 2B), such as stroma and cancer content in Factors 1,3 and 5, ISUP grade in Factors 1 and 3. Features associated with PCa aggressiveness, castrate resistance and metastasis status were captured in Factors 1,3, 4 and 5. We chose to further explore the first 5 factors because they explained the most variance (Figure 2C) and their strong correlations with clinical features (Figure 2B, Supplementary Figure 2).

To gain an understanding of the biological mechanisms underlying each factor, we performed GSEA separately on genes with positive and negative weights in each factor (Figure 2D). GSEA revealed seven biological themes that emerged as sources of variation based on the first five factors (Figure 2D). As this analysis focuses on exploring the relationship between various omics, we excluded the biological themes “lipid oxidation” and “pH regulation” due to a lack of significant correlations (Spearman’s ρ > 0.45) with either metabolomics or epigenomics measurements, resulting in five biological themes for further exploration (Figure 2D): Response to zinc ion, smooth muscle, cell cycle, immune response, and tissue morphogenesis.

To further explore biological themes, we focused on genes and metabolites with the highest absolute weights in factors associated with each theme (Figure 2D, Supplementary Figure 3). For DNA methylation, we selected CpGs with positive weights in the highest quartile and negative weights in the lowest quartile to perform enrichment analysis for chromatin states (from PCa cell lines LNCaP and PC3) and transcription factor binding regions, to assess whether CpGs are involved in regulatory areas of the genome.

**Figure 3:**
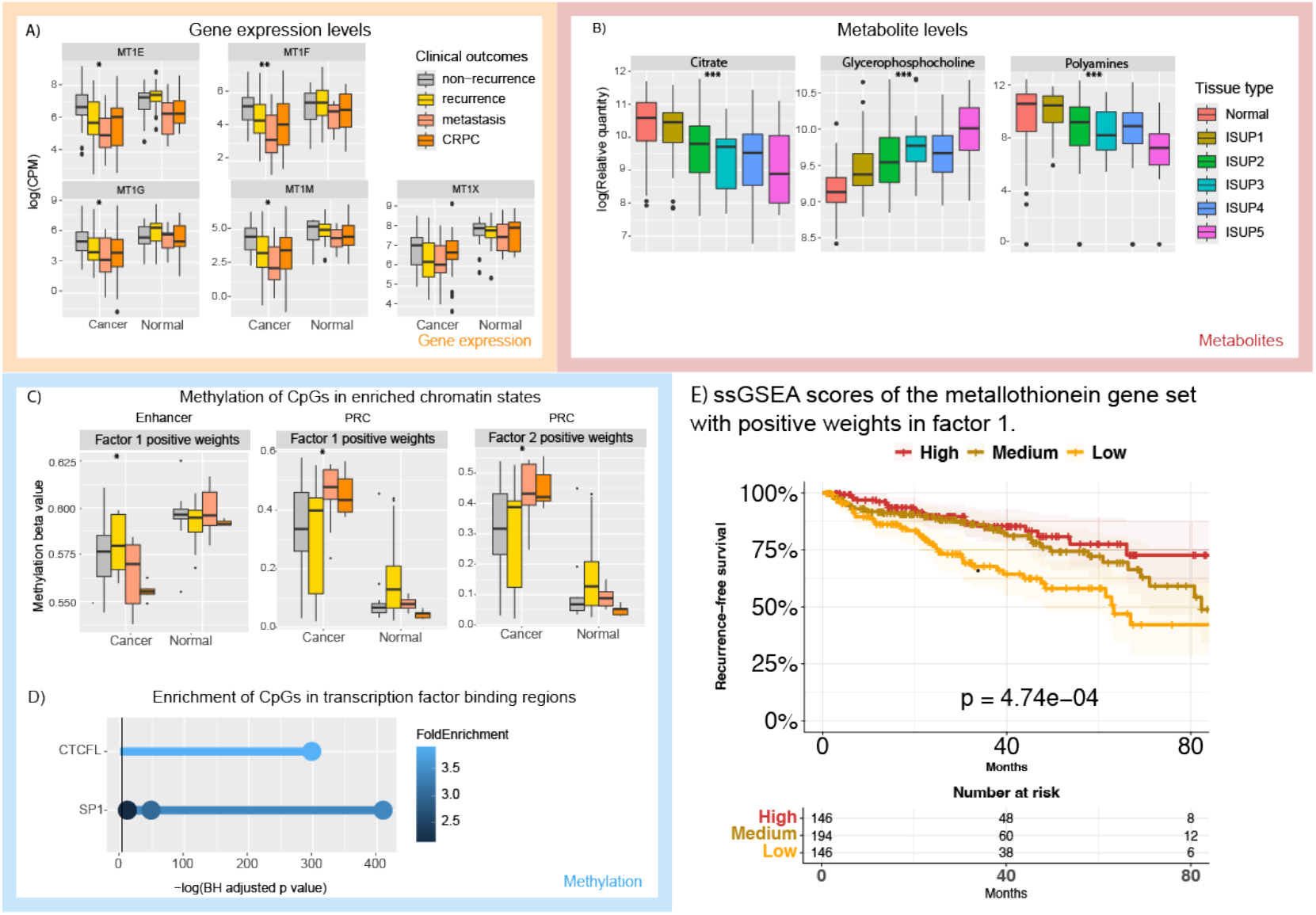
Response to zinc ion. **A)** The metallothionein gene family (MT1E, MT1F, MT1G, MT1M, MT1X) shows a trend of decreasing expression in samples with recurrence, metastasis and castrate resistance (CRPC) compared to non-recurrence samples. **B)** Citrate and polyamine levels, which have high positive weights in Factors 1 and 2, also show a trend of reduced levels in higher ISUP grades. While glycerophosphocholine shows increasing levels with ISUP grades. **C)** Methylation of CpGs in enhancers shows increased methylation in cancer samples compared to normal, and a trend towards higher methylation in recurrence, metastasis and CRPC compared to non-recurrence (Factor 2 positive weights). The same pattern is observed in polycomb repressive complex (PRC) for Factor 2 positive weights. **D)** CpGs with high positive weights in Factors 1, 2, and 5 are enriched in transcription factor binding regions of CTCFL and SP1. **E)** The ssGSEA scores of the metallothionein gene set were significantly different when separating into high, low and medium risk for recurrence-free survival in the TCGA validation cohort. * = P-value < 0.5; ** = P-value < 0.1; *** = P-value < 0.01

### Loss of metallothionein expression, citrate and polyamine levels are associated with loss of normal prostatic functions in cancer development

In Factors 1, 2 and 5, genes that were associated with the gene ontology term “the response to zinc ion” were enriched (Figure 3). The metallothionein gene family (MT1E, MT1F, MT1G, MT1M, MT1X), an important gene family in the “response to zinc ion” GO term, showed a trend of decreasing expression in samples with recurrence, metastasis and castrate resistance (CRPC) compared to non-recurrence samples both in cancer and normal tissue samples (Figure 3A). Metallothioneins can bind and transport zinc, an important metal in the normal prostate (30). High metallothionein levels lead to zinc accumulation, which further inhibits the TCA cycle, causing an accumulation of zinc and citrate in normal prostate cells, which is then secreted as key components of the prostatic fluid (31). Together with another prostatic fluid component, the polyamines, citrate and the polyamines showed a trend of lower levels in samples with increasing ISUP scores in Figure 3B. This coincides with what has been thoroughly studied and presented in previous metabolomics studies by our group (14). The lower expression of metallothionein genes may be the cause of less zinc transported into the mitochondria that again lead to higher consumption of citrate due to less inhibition of the TCA cycle (31). In previous studies, we have associated lower zinc levels with higher ISUP grades (31) and lower citrate levels (32). Both reduced citrate and the polyamine levels, which we found to be highly correlated (33), have previously been connected to aggressive prostate cancer and recurrence (16).

The CpGs with positive weights in Factors 1,2 and 5 were enriched for enhancer and PRC chromatin (Supplementary Figure 4) and with transcription factor binding regions of CTCFL and SP1 (Figure 3C, 3D). Enhancer CpGs in F1 with positive weights showed lower methylation in cancer compared to normal, while CpGs in PRC regions in F1 and F2 showed increased methylation in cancer and in more aggressive outcomes (Figure 3C). Enhancer methylation is often associated with gene repression(34), suggesting that reduced metallothionein expression may be partly due to epigenetic regulation. SP1 is linked to prostate cancer aggression in several ways through both proliferation (35) and recently through induced EMT in CTCs of prostate cancer (36, 37). CTCFL is a paralog of the chromatin architectural protein CTCF and has been shown to aberrantly reorganize chromatin and to regulate AR in prostate and ovarian cancer (38).

**Figure 4:**
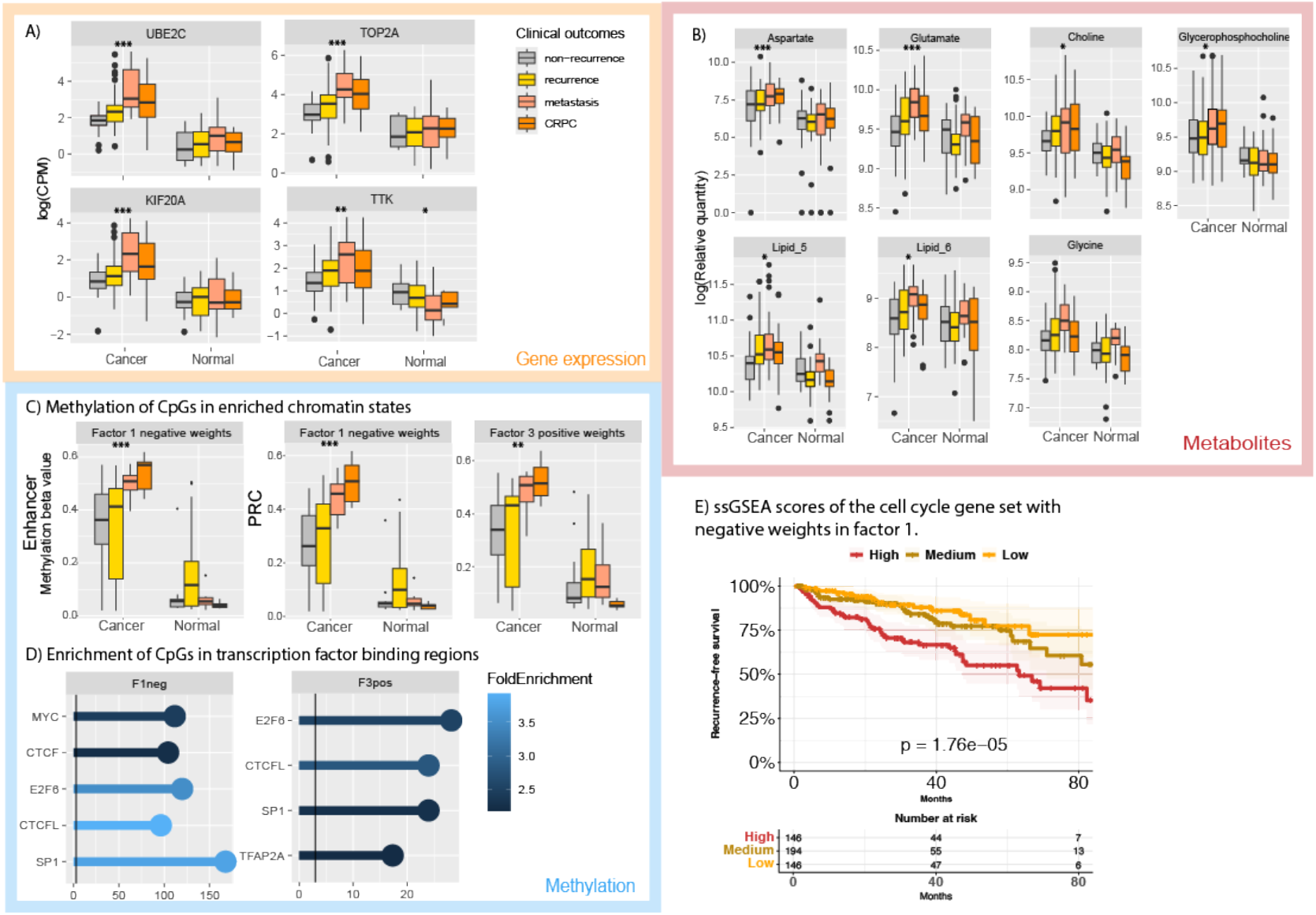
Cell cycle. **A)** Gene expression shows a trend of higher expression in cancerous samples compared to samples with non-cancerous tissue. **B)** Metabolites related to lipid and choline metabolism show a strong correlation with genes associated with cell cycle progression. **C)** The CpGs were enriched for polycomb repressive complex (PRC) and enhancer chromatin states (Supplementary Figure 4), exhibiting a higher methylation in cancerous samples than normal samples. **D)** CpGs were also enriched in transcription factor binding regions associated with cell proliferation. **E)** Validation of gene signature based on MOFA factor weights. * = P-value < 0.5; ** = P-value < 0.1; *** P-value < 0.01

To validate our gene expression metallothionein findings, we separated the ssGSEA scores of the metallothionein gene set into high, low and medium risk for recurrence-free survival in the TCGA validation cohort (Figure 3E, Supplementary Figures 5,6). Using 658 patients with up to 80 months of follow up, we found highly significant separation between the three risk groups (p=0.0004). To conclude, we found and validated decreasing metallothionein expression and higher methylation with prostate cancers aggressiveness, while citrate and polyamines levels decreased with higher sample ISUP grades. We hypothesize that metallothionein expression influences citrate levels through zinc binding and that metallothionein expression might be regulated by methylation and chromatin reorganization. These results demonstrate how metabolism, gene expression, and methylation might be interconnected in regulating citrate and zinc metabolism during the transition from normal prostate epithelial function to cancer development.

**Figure 5:**
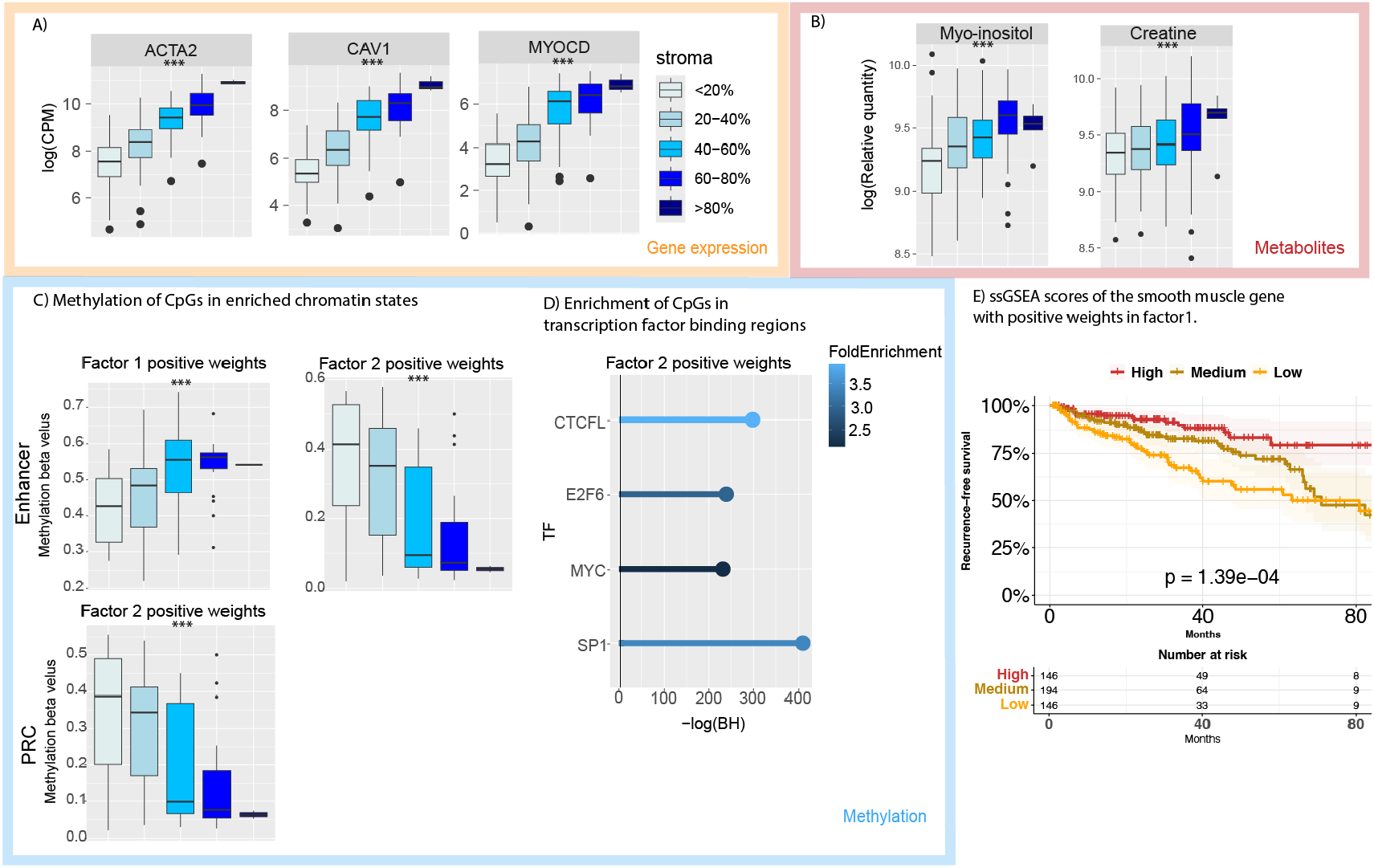
Smooth muscle. **A)** Expression levels of positively weighted genes from Factors 1 and 2, including stromal and smooth muscle-associated genes (ACTA2, CAV1, and MYOCD) and immune related CD38 gene, across samples with varying stromal proportions. **B)** Abundance of stromal-related metabolites (Myo-inositol and Creatine) in relation to tissue stromal content. Color code follows from A. **C)** Methylation of CpGs in chromatin states enriched for enhancer and polycomb repressive complex (PRC). An inverse trend of enrichment with stromal content was observed between Factors 1 and 2. Color code follows from A. **D)** Enrichment of CpGs in transcription factor binding regions, highlighting associations with CTCF, E2FC, MYC, and SP1 binding sites for positive weights in Factor 2. **E)** Single-sample gene set enrichment analysis (ssGSEA) of smooth muscle-related genes in the TCGA prostate cancer cohort, showing lower recurrence-free survival among patients with low smooth muscle gene expression scores. * = P-value < 0.5; ** = P-value < 0.1; *** P-value < 0.01

**Figure 6:**
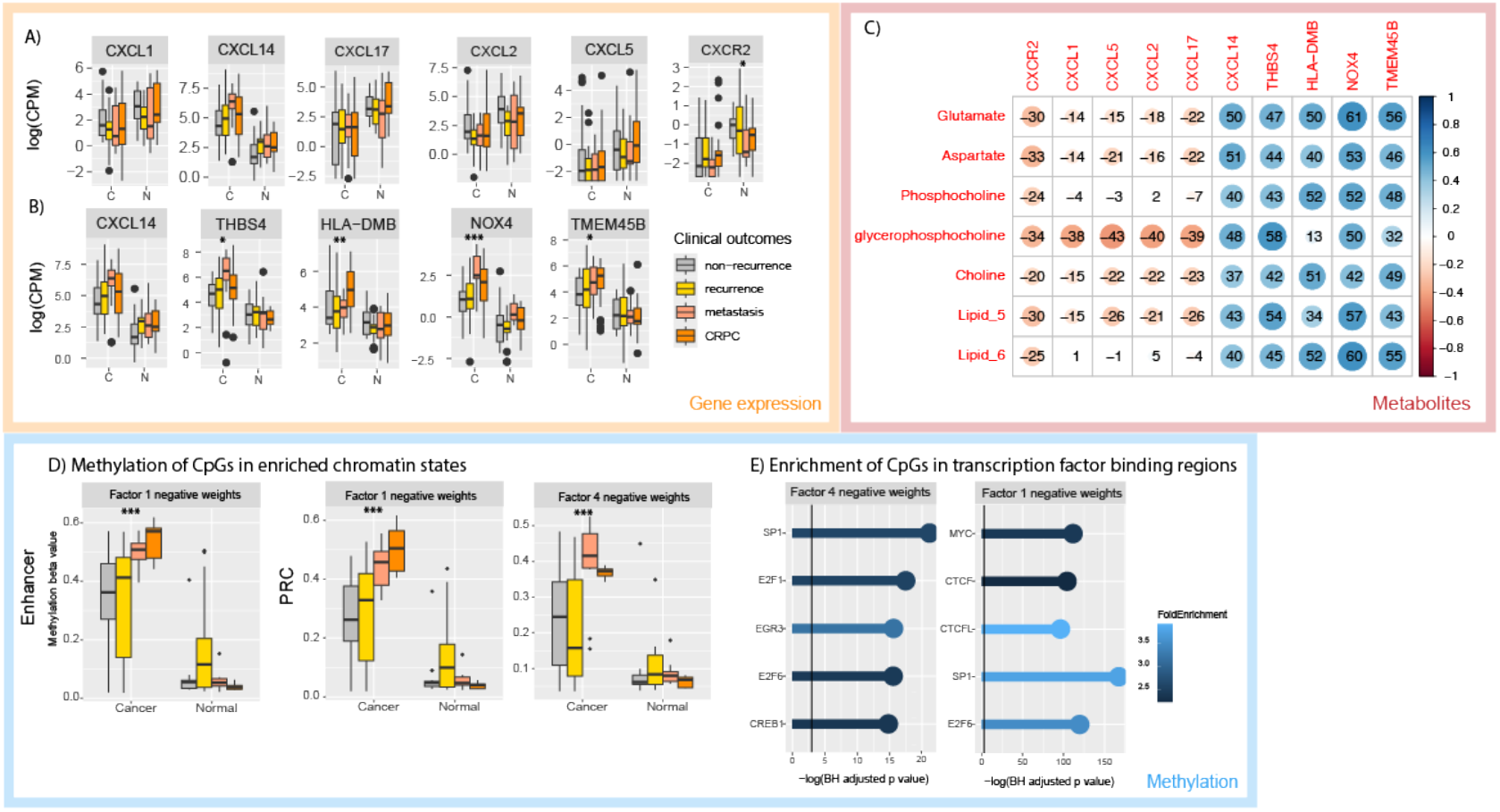
Immune activation. **A)** Gene with strong weights with elevated expression in normal (N) samples is shown with their association with cancer aggressiveness. The immune activation genes upregulated in normal samples included several C-X-C motif chemokine ligands (CXCL). **B)** Genes with strong weights with elevated expression in cancer (C) samples are shown with their association with cancer aggressiveness. **C)** Correlation between genes with strong weights from A) and B) with metabolite levels. The immune-related genes associated with cancer samples are positively correlated with metabolic glutamate levels and changes to lipid metabolism citrate and polyamine levels. **D)** Methylation sites with strong weights in Factors 1 and 4 and their overlap with chromatin states characterizing enhancers and polycomb repression (PRC), suggesting that epigenetic changes can be a driver for immune related changes in cancer development. **E)** Methylation sites with strong weights in Factors 1, 2 and 4 are enriched for binding sites for transcription factors involved in chromatin remodeling and proliferation. * = P-value < 0.5; ** = P-value < 0.1; *** P-value < 0.01

### The mitotic spindle checkpoint is associated with sample cancer content and lipid metabolism

GO terms related to cell cycle were enriched with negative weights in Factors 1 and 2, as well as positive weights in Factors 3 and 5, though only genes from Factors 1 and 2 were remaining after quantile filtering. These genes exhibit increased expression in cancer samples compared to normal samples (Figure 4A) and are associated with various components of the mitotic spindle, including the centromere (TTK) and the anaphase-promoting complex (KIF20A, UBE2C, and TOP2A)(39, 40). TDRD1 has been identified as a direct target gene of ERG overexpression in primary prostate cancer. The mitotic spindle checkpoint ensures that cells do not divide until all chromosomes are properly aligned. In cancer, this checkpoint may become disrupted, allowing cells to proceed with division even if chromosomes are not correctly aligned (41, 42). This disruption may lead to accelerated cell cycle progression, chromosomal instability, and aneuploidy, all of which are commonly associated with cancer in general as well as prostate cancer (43, 44).

The genes further show a strong positive correlation with several metabolites which have negative weights in Factor 1 and positive weights in Factors 3 and 5 (Supplementary Figure 7) All of these metabolites had higher levels in cancer samples compared to normal (Figure 4B). Glutamate, aspartate, and glycine are involved in glycolysis and augmented amino acid metabolism, which are tightly coupled with an increased flux of the TCA cycle. They offer building blocks and energy for rapid cell proliferation (45). Choline functions as an important methylation donor that can affect DNA methylation (46) and lead to disruption of DNA repair (47). Further, choline, glycerophosphocholine, and lipids are essential for cell membrane synthesis and phospholipid metabolism, linking these metabolites to the cell cycle and increased cell turnover (48). Elevated levels of phosphocholine and increased cell turnover is a well-known feature of prostate cancer (49).

**Figure 7:**
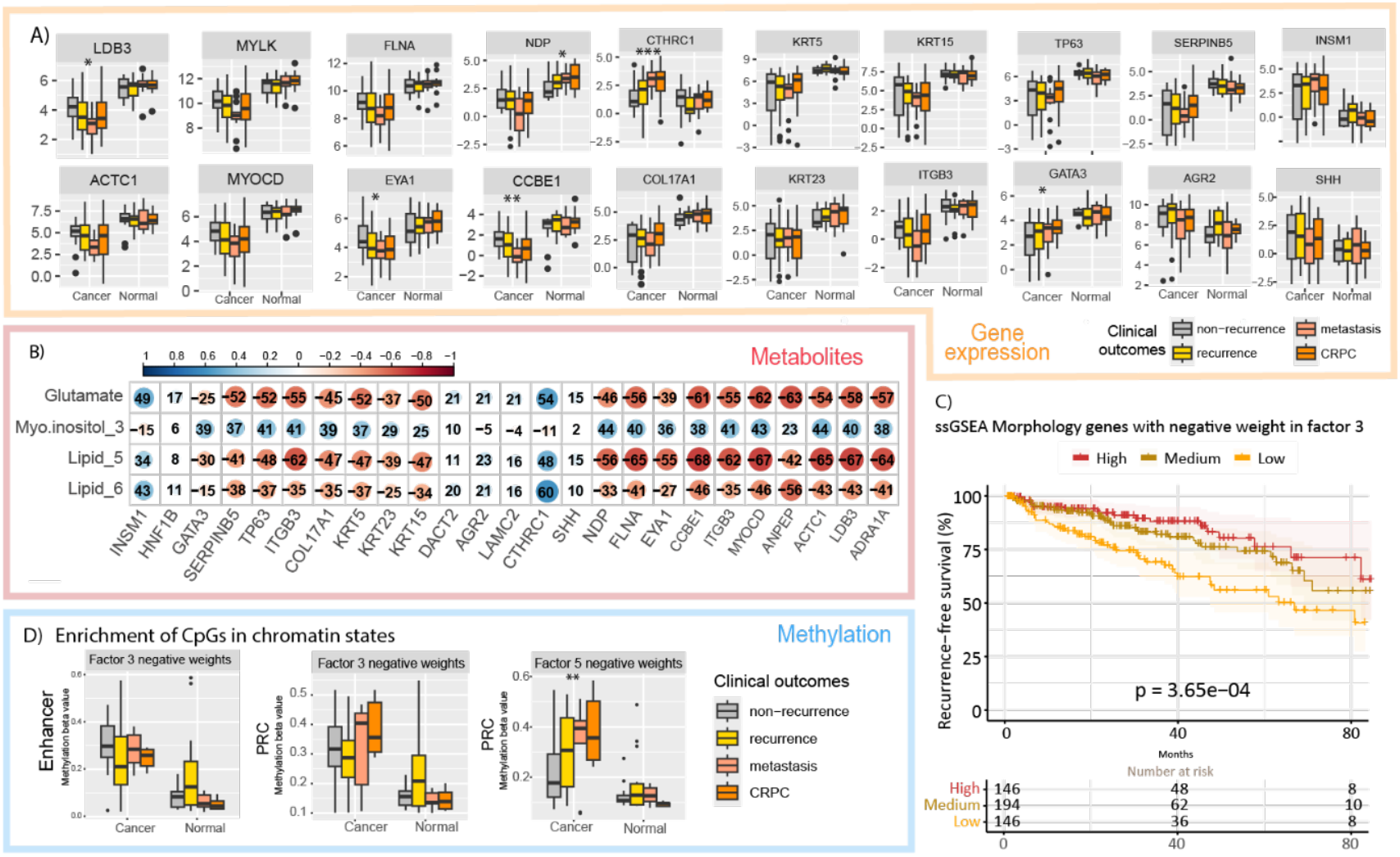
Tissue morphology. **A)** Gene expression of morphology genes related to stroma and glandular morphology with negative feature weights in Factors 3, 4, and 5. **B)** Correlation plot (Spearman) between morphology genes and metabolites. **C)** Kaplan-Meier survival analysis based on GSEA with the morphology genes with negative feature weights for Factor 3. **D)** Enrichment of CpGs in chromatin states across negative Factors 3 and 5. *\* = P-value < 0*.*5; ** = Pvalue < 0*.*1; *** = P-value < 0*.*01*

In addition, the genes are strongly correlated (Spearman’s ρ > 0.5) with several CpGs (Supplementary Figure 4). These are enriched in polycomb repressive complex (PRC) and enhancer regions (Figure 4C) which showed increased methylation in samples with cancer compared to those without cancer, and highest methylation in samples with aggressive disease. Further, the strongly correlated CpGs were also enriched in binding regions of several TFs (Figure 4D), including MYC, CTCF, E2F6, CTCFL, and SP1, which are known to regulate proliferation of prostate cancer cells (50, 51).

We validated the cell cycle related genes as a gene signature and found a significant separation between the three risk groups (Figure 4E; Supplementary Figure 8).

The metabolites, genes, and methylation patterns associated with the cell cycle theme are not directly connected through a single pathway but are all essential for cell division. Gene expression regulates the mitotic spindle checkpoint, metabolism synthesizes cell membrane, and methylation rearranges chromatin to allow for cell cycle progression. Together, these processes promote cell proliferation.

### Smooth muscle-specific omics characteristics correlate with stromal enrichment

The top positively weighted genes for the “smooth muscle” theme in Factors 1 and 2 included ACTA2 and MYOCD, which are canonical markers and regulators of smooth muscle differentiation, along with CAV1, a gene broadly expressed in stromal and vascular cells. Interestingly, expression levels of ACTA2, CAV1, and MYOCD increase progressively with higher stromal content. ACTA2 is primarily expressed in the smooth muscle cells of the prostate stroma, and downregulation of ACTA2 expression has been linked to poorer overall survival in PCa patients (52). Similarly, loss of CAV1 in the stroma is a strong marker for increased prostate cancer progression (53, 54). Although MYOCD has not been explicitly studied in the context of PCa, it is recognized as a tumor suppressor in lung cancer (55). The observed higher expression of stromal genes in regions with abundant stroma suggests that Factors 1 and 2 may reflect normal prostate stromal characteristics. Moreover, the observed higher expression of smooth muscle genes mirrored the higher levels of metabolites such as myo-inositol and creatine in samples with a higher stroma content (Figure 5A and 5B). We have previously detected creatine at higher levels in stroma with spatially resolved MSI (32). Creatine plays a vital role in muscle cells by regenerating ATP through the phosphocreatine–creatine kinase system and thereby maintaining ATP homeostasis during high-energy situations (56). The exogenous myo-inositol functions as an epithelial-mesenchymal transition (EMT) inhibitor (57), and increased levels in stroma-rich samples may suggest a local stromal defense.

Factor 1 and Factor 2 captured distinct CpG methylation patterns within enriched chromatin states, where Factor 1 showed an increase and Factor 2 a decrease in enhancer methylation beta values with increasing stromal content. This highlights two opposite stromal-associated epigenetic re-programming patterns (Figure 5C). High-weight probes from both Factors 1 and 2 are provided in the Supplementary Figure 4. The CpGs were enriched in transcription factor binding sites (including CTCFL, E2F6, MYC, and SP1), pointing to regulatory mechanisms such as chromatin remodeling, oncogenic transcriptional activation and stromal remodeling programs (Figure 5D). Importantly, analysis of the TCGA cohort showed that reduced activity of a smooth muscle gene program correlated with significantly worse recurrence-free survival also in the TCGA cohort, suggesting that preservation of smooth muscle-like stromal characteristics may be associated with a more favorable prognosis (Figure 5E; Supplementary Figure 9).

### Different sets of genes are associated with immune activation in cancer and normal samples, and are related to DNA methylation changes (in enhancer and Polycomb genomic regions)

The terms related to immune activation are enriched in Factors 1, 2 and 4 (Figure 2D), with Factor 1 and 4 somewhat stronger than Factor 2. In addition to RNA and metabolites, these three factors are characterized by being strongly influenced by changes in DNA methylation (Figure 2C). There is a slight overall association with ISUP grade and delated clinical variables (clinical t-stage, seminal vesicle invasion, recurrence status, metastasis status and castrate resistance, Figure 2B), but for sample characteristics there is no overall strong association with cancer and stroma content.

We identified two subsets of immune-related genes associated with the three immune activation factors, where one subset was more enriched in cancer samples and one subset was more enriched in normal samples. The subset enriched in normal samples included several C-X-C motif chemokine ligands (CXCL) (Figure 6A). Interestingly, three of these chemokines (CXCL17, CXCL2 and CXCL5) are part of a gene signature recently published by our research group which characterize local immune modulation and invasion of club-cells in normal appearing glands adjacent to aggressive tumors (58). This demonstrates that the MOFA-factorization can capture trends in local tissue compartments which otherwise could only be identified by more advanced methods such as Spatial Transcriptomics. The immune-related genes that MOFA associated with cancer samples were positively correlated with glutamate levels and changes in lipid levels (Figure 6C).

We observed a strong association of Factors 1, 2 and 4 with changes in DNA Methylation. Methylation site changes with high absolute weights in these factors are enriched in enhancers and polycomb (PRC) regulatory regions, and these changes are more prominent in aggressive cancer samples (Figure 6D). This suggests that the epigenetic changes can be driven by immune-related processes in cancer development. These methylation sites were also enriched for transcription factors known to be involved in chromatin remodeling (CTCF and CTCFL)(38, 59) and tumor proliferation (MYC, E2F, SP1) (Figure 6E) (51, 60-62). These changes can be caused both by active chromatin remodeling of prostate epithelial cells or by infiltration of foreign cell-types (for example Club cells) in the tissue environment. However, changes in polycomb-regulation is generally associated with developmental programs (63).

Although the Kaplan-Meier analysis showed no significant outcome for patients with intermediate levels of immune related genes (Supplementary Figure 10-12), our immune activation results may indicate that balanced immune activation is more beneficial for tumor development (64).

### Stromal and glandular morphology gene features show an inverse association to cancer characteristics

GO terms related to morphology and tissue development (n = 8) all showed a similar pattern across Factor 3, 4 and 5 with negative feature weights (Figure 2D). Tissue morphology refers to the structural organization of tissue. In prostate, the key tissue compartments are the glandular structures and the surrounding stroma. Factor 3 correlated strongly with both stroma content and cancer content, though in opposite directions (Figure 2B), whereas these relationships were much weaker for Factors 4 and 5.

Four of the eight enriched GO terms were related to gland morphology and one to appendage development, while three GO-terms gave the indication of enriched vasculature function. However, further inspection of the genes within the vasculature GO-terms and interpreting them in the context of prostate tissue biology, indicated that this likely represents enrichment of stroma-related gene expression, rather than vasculature morphology. This is showcased by enrichment of MYLK, MYOCD, ACTC1, LDB3 and FLNA which are all key genes of muscle fibers of smooth muscle cells (65). These genes had the strongest negative factor weights for Factor 3 (compared to Factors 4 and 5) and systematically had the highest expression in normal prostate samples (Figure 7a). Further, all these stroma genes had a high inverse correlation (r => -0.55) with Lipid_5 (Figure 7b). As previously presented, Lipid_5 showed higher levels in cancer samples compared to normal samples (Figure 4B). Although we do not have the precise identity of the lipids contributing to this NMR-signal (ppm-range 3.68-3.79), higher lipid concentration is a well-known characteristic of prostate cancer (49). It is therefore expected that stroma-genes associated with healthy tissue have an inverse relationship with lipids.

Exploring the genes within gland-morphology GO terms showed that most were linked to basal cells, which help maintain normal prostate gland integrity. Proteins that contribute to this physical integrity make up the intermediate filaments and include cytokeratin genes such as KRT5, KRT15 and KRT23 which had strong negative feature weights for both Factors 3 and 5 and were downregulated in cancer samples (Figure 7a). As loss of basal cells is one of the first indications of cancer, cytokeratin 5 (KRT5) is routinely used in the clinic to confirm the presence of basal cells and to differentiate benign glands from invasive carcinoma (66). Another clinically used marker for basal cells that had high negative feature weights (Factors 3 and 5) is TP63 (Figure 7A). TP63 (p63) is a transcription factor and a known tumor suppressor of prostate cancer (67). In line with previous scientific consensus, TP63 had reduced expression in cancer samples (Figure 7A) (67). GATA3 is also a transcription factor with tumor suppressive properties that showed the same pattern (68). Other genes related to basal cells and basement membrane with negative feature weights in Factors 3 and 5 included SERPINB5, ITGB3 and COL17A. TP63, ITGB3 and SERPINB5 had a strong inverse correlation with glutamate (r ≥ -0.52, Figure 7B). As previously mentioned, glutamate was found at higher levels in cancer compared to normal tissue both in this and other studies (45). Similarly to the stroma-related genes, the gland genes were also inversely correlated with Lipid_5.

Negative Factor 3 is capturing features related to normal prostate tissue morphology, indicated by genes known to be present in healthy stroma and glands had negative weights and by the stroma content pulling in the same direction (Figure 2B). On the other hand, cancer content and recurrence status had an inverse relationship with negative factor 3 (Figure 2B). This inverse association to clinical outcome was validated in the TCGA-cohort, where a high ssGSEA score for morphology-genes (with high negative feature weights in factor 3) was protective against recurrence (p = 0.0004, Figure 7C, Supplementary Figures 13-14).

Some genes within this morphology theme only had negative feature weights for Factor 5 and generally were less prominent in Factors 3 and 4 (Supplementary Table 1). Interestingly, several of these genes, including INSM1, SHH and AGR2 had an increased expression in cancer samples compared to normal (Figure 7a). These genes have more diverse functions with INSM1 being a transcription factor (69), while AGR2 is involved in protein folding (70) and SHH is important prostate development and regeneration (71). Factor 5 is clearly capturing other morphology functions that are not highly associated with neither normal or cancer morphology but rather represents variation that is independent of stroma and cancer content.

The identified CpGs with negative weights in Factors 3 and 5 were enriched in enhancers and PRC regions (Supplementary Figure 4) and were associated with clinical outcomes (Figure 7D). The CpG enrichments in enhancers and PRC regions were in line with the other factors and biological themes, suggesting that chromatin remodeling plays a part in several cancer driving biological functions of prostate cancer.

## Discussion and concluding remarks

Our multi-omics analysis of both normal and cancerous prostate tissues has unveiled several distinct biological themes associated with prostate cancer and prostate cancer aggressiveness. The advantage of employing MOFA in this study lies in its ability to integrate heterogeneous data modalities while robustly handling missing observations across datasets. Although this linear framework may not capture complex non-linear relationships, its high interpretability proved valuable for elucidating the connections between molecular features and prostate cancer biology. Consistent with the model’s architecture, Factor 1 accounted for the greatest proportion of variance across all three omics layers and exhibited the strongest correlations with clinical characteristics (Figure 2A, B). This suggests that the dominant source of variation in the dataset is inherently linked to cancer biology and disease progression. Furthermore, the highest enriched GOs were found in Factor 1 and were related to cell cycle (Figure 2D). Dysregulation of the cell cycle is a particularly well-known hallmark of cancer (43, 44), and these findings therefore prove that MOFA is accurately capturing the important variation associated with cancer aggression.

The precision of MOFA allowed us to link aspects of prostate cancer molecular biology that had not been previously studied across multiple omics levels. The expression of metallothioneins and their potential connection to citrate have been documented (33), but our study suggest that they might be regulated by promoter methylation. Similarly, the mitotic spindle checkpoint in prostate cancer has traditionally been examined one gene at a time, but we identified a pattern of increased expression across multiple checkpoint genes. Although cancer-related metabolism and the loss of basal cell fate are well-known phenomena (72, 73), our study suggest they might be regulated by chromatin state rearrangements. While the immune microenvironment in prostate cancer has been described in single-cell or spatial contexts (74-77), our study pinpoints chemokines and immunoglobulins as important elements of the prostate cancer tumor microenvironment.

Factors 1 and 3 captured molecular variations associated with ISUP grade, cancer content, and stroma content, which are challenging to disentangle using bulk data. Prostate cancer originates from epithelial cells and will therefore be a source of epithelial genes and regulatory mechanisms. Stroma on the other hand, is displaced by growing tumors, and thus a “normal” sample also includes more stromal tissue than one with cancer, leading to imbalanced stroma signals.

In addition, samples with higher ISUP grade often have a larger cancer area and thus less stroma content introducing a bias in the analyses. Due to this, separating biological mechanisms coming from ISUP grade, cancer content and stroma is not feasible with the current experimental design. A clear next step is to explore the biological patterns from this study further using spatial technologies and/or dissected tissue compartments.

Our integrative MOFA analysis revealed that CpGs contributing to multiple molecular factors in prostate cancer are enriched both in Polycomb repressive complex (PRC)-associated chromatin states (Figures 3-7) and in binding regions of the transcription factors SP1 and CTCFL (BORIS). This enrichment points to an epigenetic landscape in which developmental regulatory elements, normally repressed by Polycomb, become accessible and engaged by transcriptional activators (78). CTCFL is known to compete for binding sites with CTCF (79) and may potentially displace PRC2 and facilitate chromatin opening (80). SP1 can occupy CpG-rich promoters upon chromatin opening to initiate transcriptional activation (81). Together, these mechanisms suggest a coordinated reactivation of Polycomb-silenced gene programs during prostate tumorigenesis.

In our analyses, SP1 and CTCFL emerge as recurrent transcriptional regulators shaping diverse biological processes captured by the MOFA factors. SP1 is a well-characterized factor in prostate cancer, promoting androgen receptor (AR) transcriptional activity (82), and its activity is often enhanced by promoter hypomethylation and increased chromatin accessibility (83). In contrast, CTCFL, a testis-specific paralog of CTCF, is only sporadically reported in prostate cancer but has been implicated in DNA hypomethylation and chromatin reorganization in other malignancies (84, 85). Its ectopic activation may alter enhancer insulation and enable germline-like gene expression. The consistent co-enrichment of SP1 and CTCFL binding regions among factor-associated CpGs across functional modules such as cell cycle, zinc metabolism, immune signaling and stromal remodeling supports a model of global epigenetic reprogramming. These findings together point to a novel interplay between Polycomb deregulation, CTCFL-mediated chromatin remodeling, and SP1-driven transcriptional activation in shaping the prostate cancer epigenome.

A limitation of this study is the unbalanced representation across omics layers (Figure 2A), as data for all modalities could not be obtained for every sample. This was due to multifactorial constraints, including limited time, resources, and the high analytical costs associated with DNA methylation and gene expression profiling. MOFA is designed to handle such missing measurements, however, extra care must be taken when interpreting the cumulative variance explained in the MOFA model. As noted by MOFA the developers, having fewer samples present in one omics dataset tend to give the more variance captured from that omics dataset (86). This is likely the case in our MOFA model, where the DNA methylation dataset (n=96) had the highest cumulative variance explained of 76% compared to the transcriptomics (50%, n=176) and metabolomics data (29%, n=280). Furthermore, the metabolic coverage was restricted to 31 metabolites via HR-MAS analyses. However, the use of this non-destructive technique was essential to enable subsequent gene expression analysis on the same physical samples. Notably, the identified metabolites represent key oncogenic pathways, including the TCA cycle, amino acid metabolism, and lipid levels.

To conclude, by integrating DNA methylation, gene expression and metabolites, we show that key prostate cancer-driving biological functions are impacted by multiple molecular alterations. Metallothionein/citrate levels, cell cycle activity, smooth muscle/stromal integrity, immune activation and tissue morphology are all functions affected non-redundantly by all three molecular layers, supporting a system-level deregulation of these key functions. The alterations in CpGs in PRC regions and binding regions of SP1 and CTCFL suggest that epigenetic reprogramming contributes to these coordinated transcriptional and metabolomic changes in prostate cancer. Gene expression signatures derived from these biological functions show validated prognostic power illustrating both biological and clinical relevance.

## Notes

### Competing Interest Statement

The authors have declared no competing interest.

### Funding

- Norwegian Cancer Society: 272770, 208263-2019, 223290
- Helse-Midt Norge: 2024-36825, 34183
- European Research Council: 758306
- Enabling Technologies, NTNU
- ELIXIR Norway: 322392

