## Supplementary Figures for "Integrated multi-omics profiling uncovers the epigenetic, transcriptional, and metabolic landscape of prostate cancer progression"

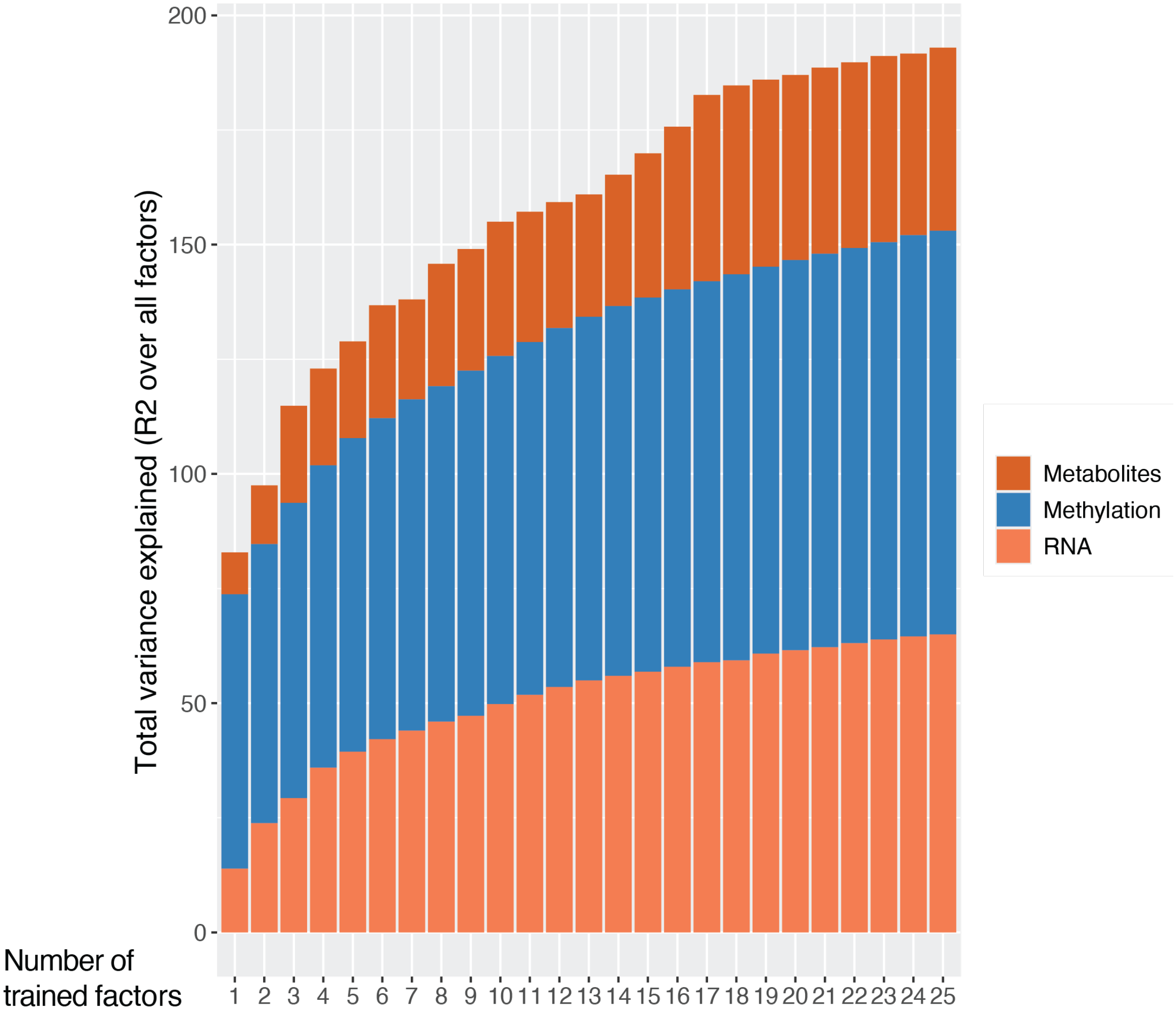


Supplementary Figure 1. Total explained variance for MOFA models with different numbers of factors. For each number of factors from 1 to 25, a separate MOFA model was trained and the total explained variance for each MOFA model was added.

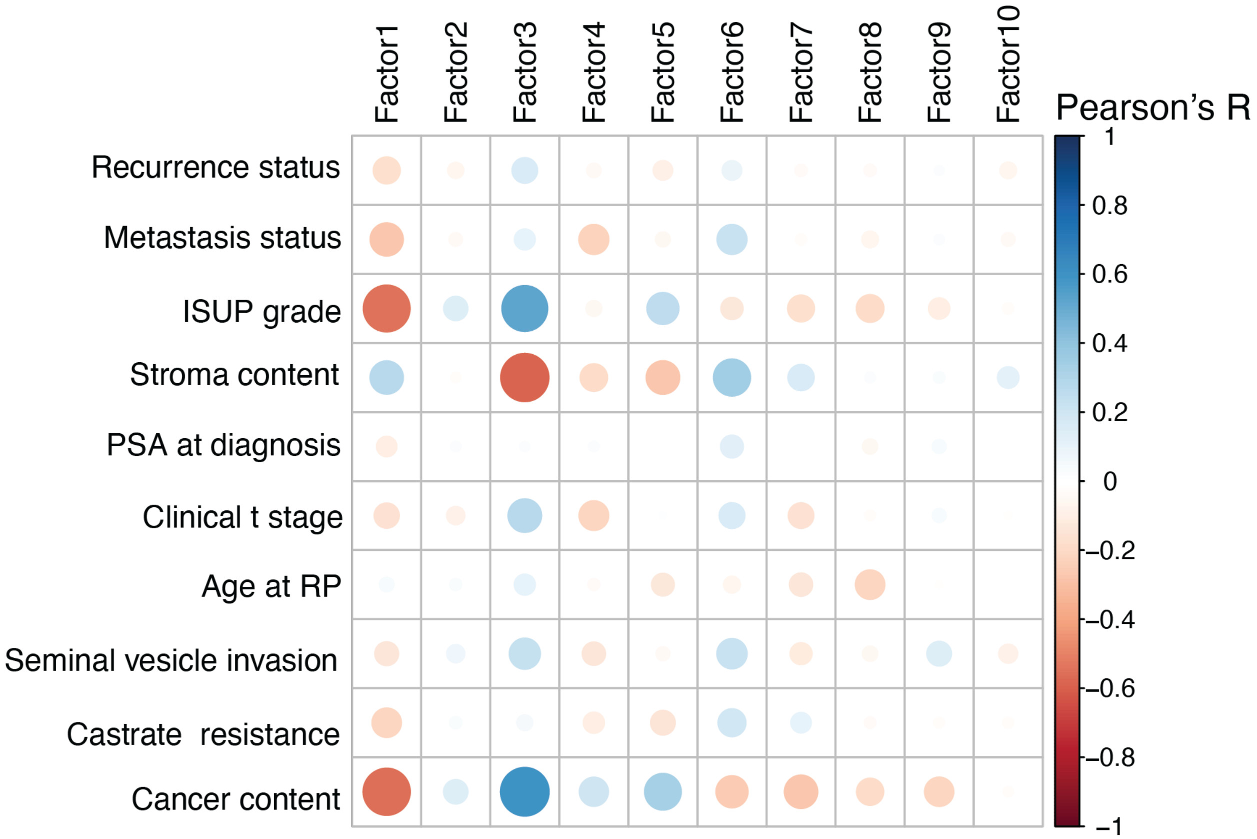


Supplementary Figure 2. Pearson correlations between all 10 factors of the final MOFA model and covariates.

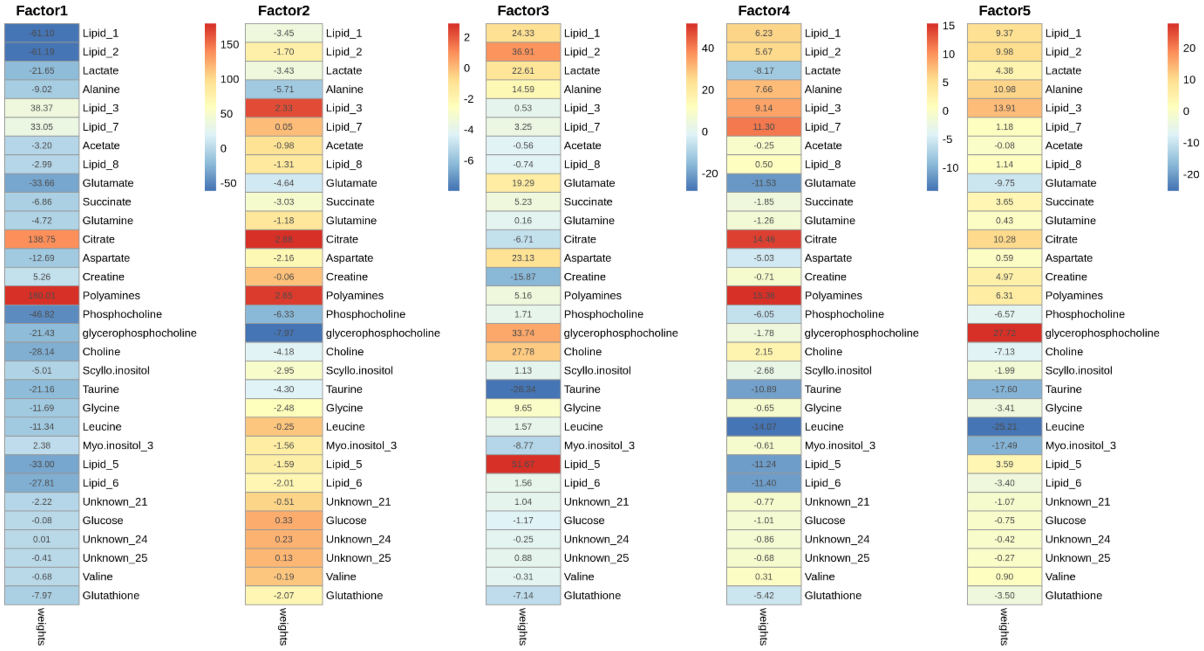


Supplementary Figure 3. Heatmap for factor weights of factors 1 to 5 for metabolite data.

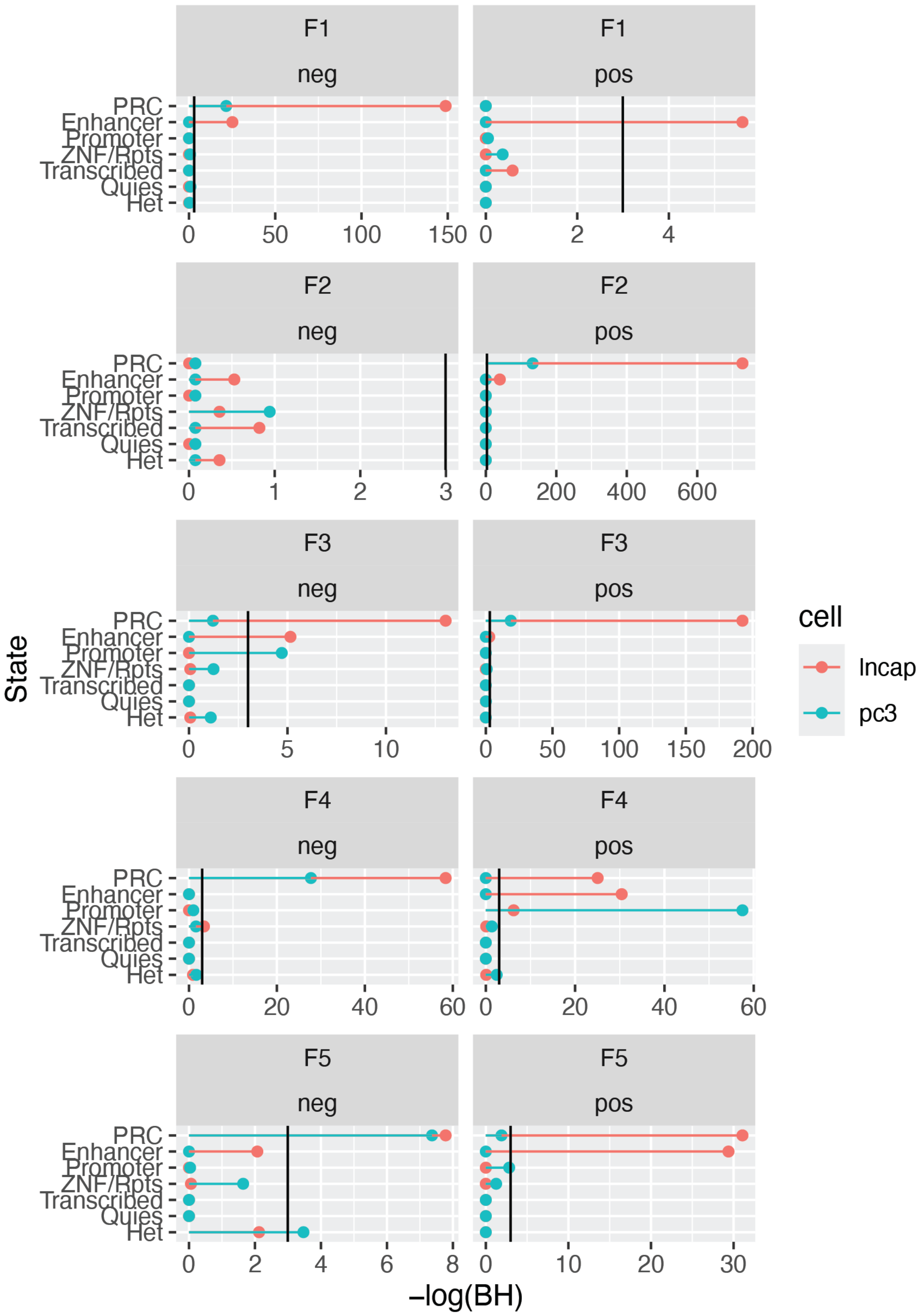


Supplementary Figure 4. Enrichment of CpGs for chromatin states for factors 1 to 5 for positive and negative factor weights.

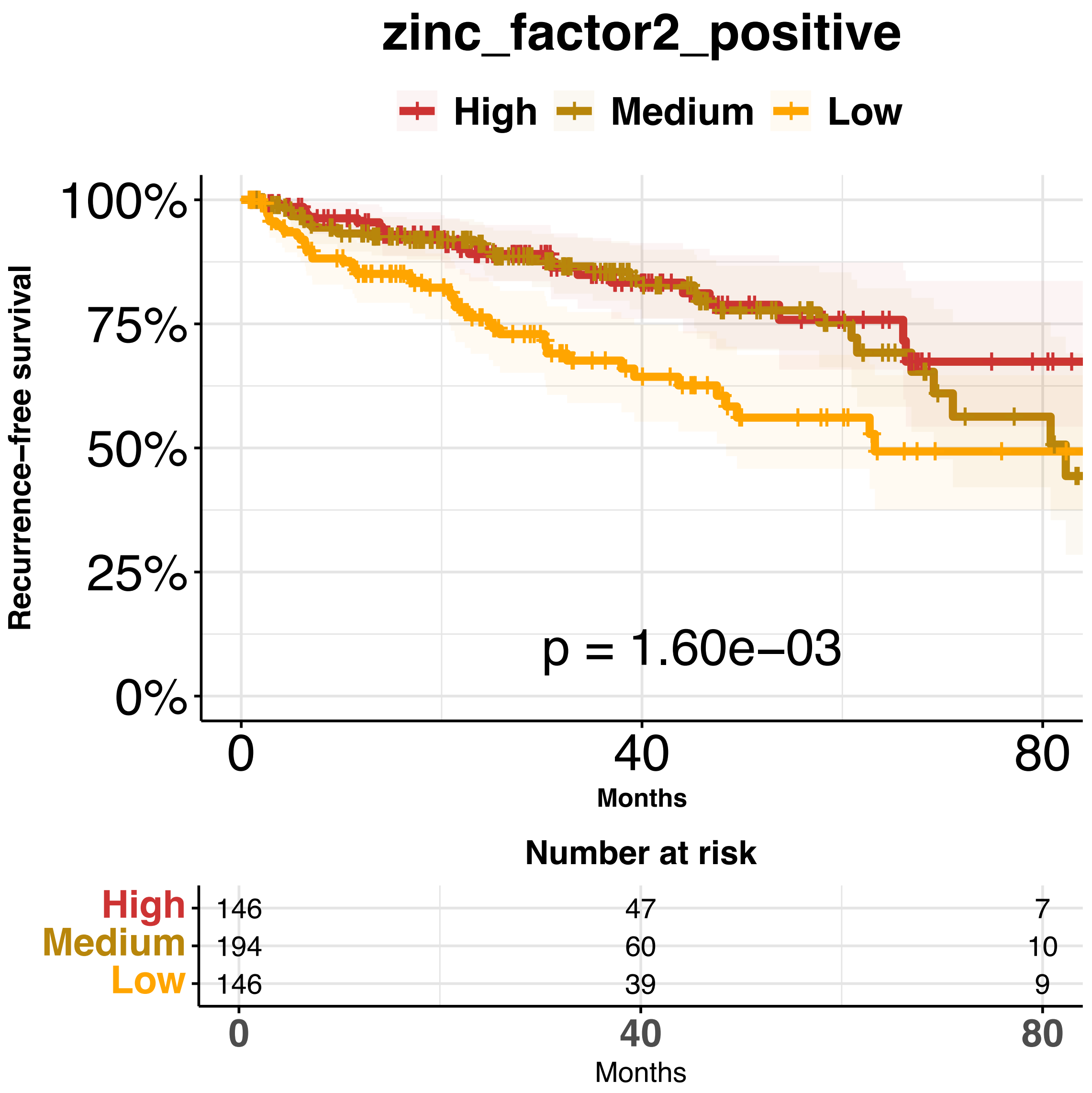


Supplementary Figure 5. Single-sample gene set enrichment for positive factor 2 weights for the biological theme “response to zinc ion”.

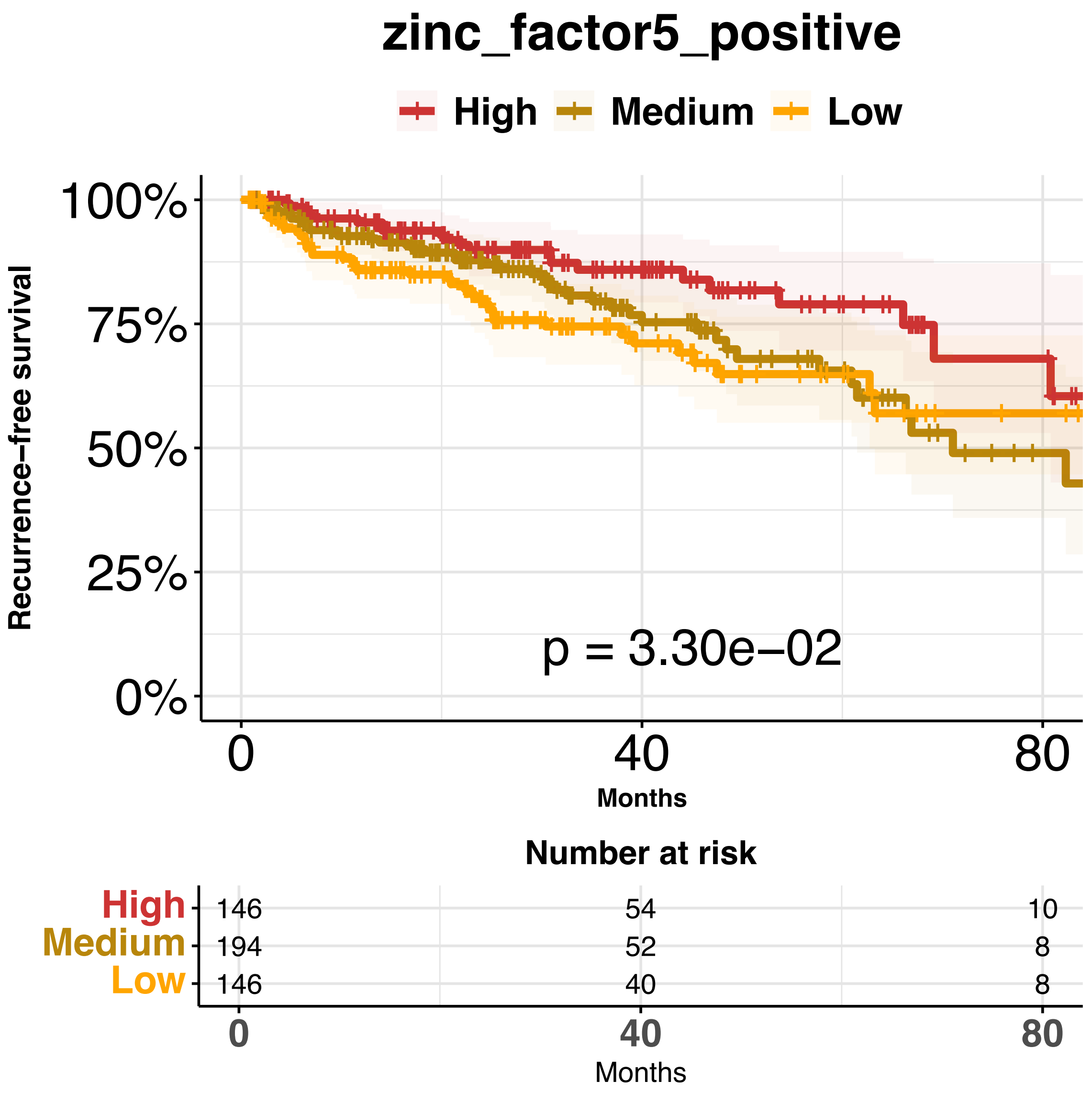


Supplementary Figure 6. Single-sample gene set enrichment for positive factor 5 weights for the biological theme “response to zinc ion”.

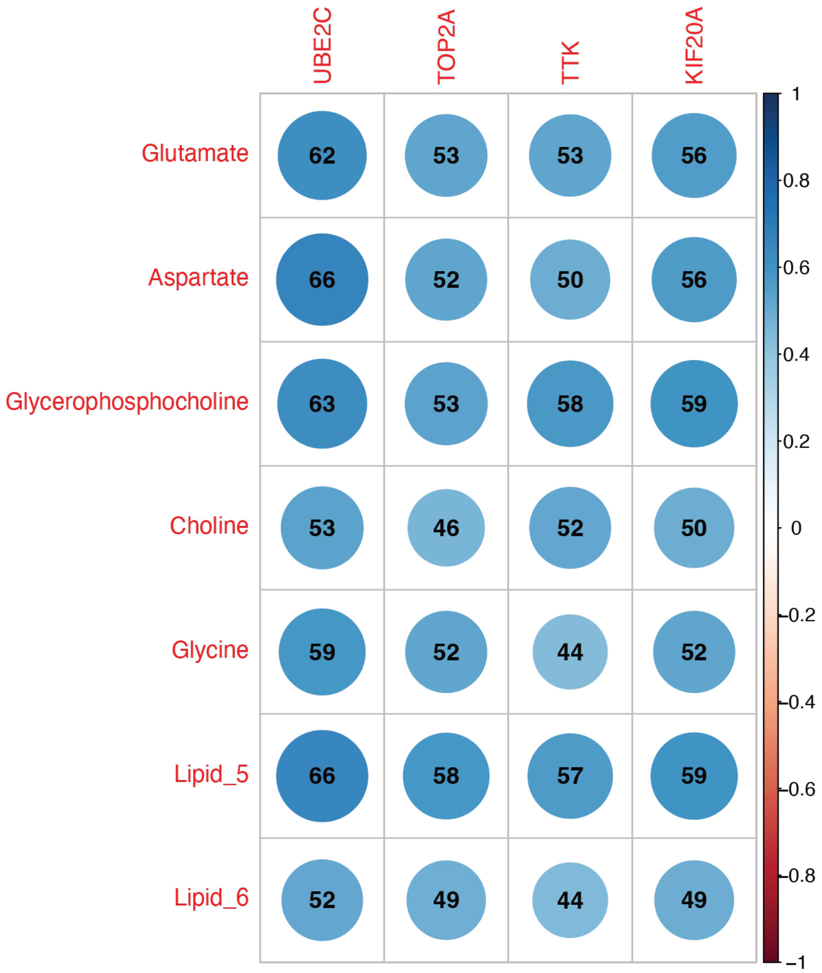


Supplementary Figure 7. Correlation between genes associated with the biological theme "cell cycle progression" and metabolites with an absolute Spearman correlation > 0.5.

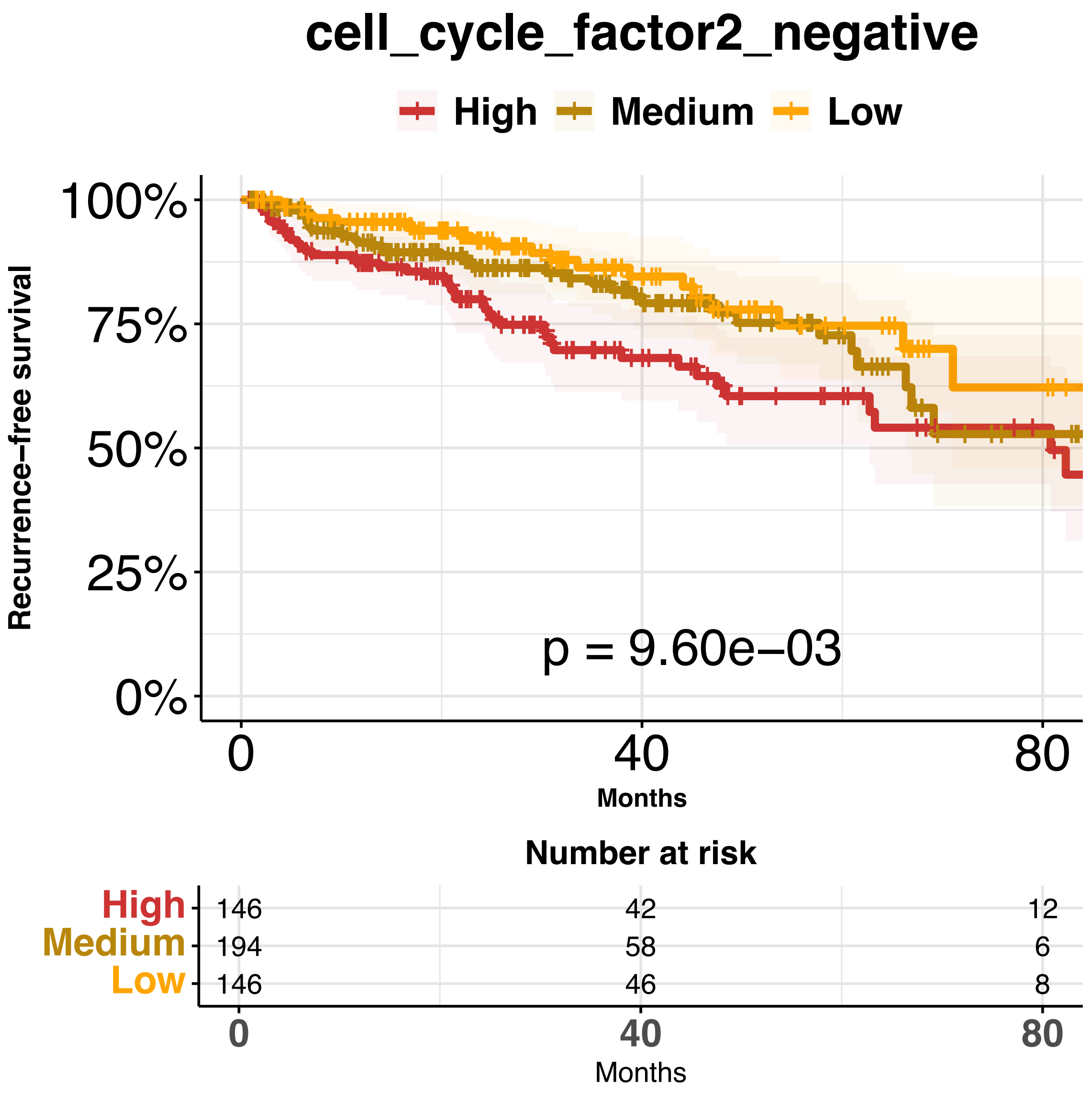


Supplementary Figure 8. Single-sample gene set enrichment for negative factor 2 weights for the biological theme “cell cycle”.

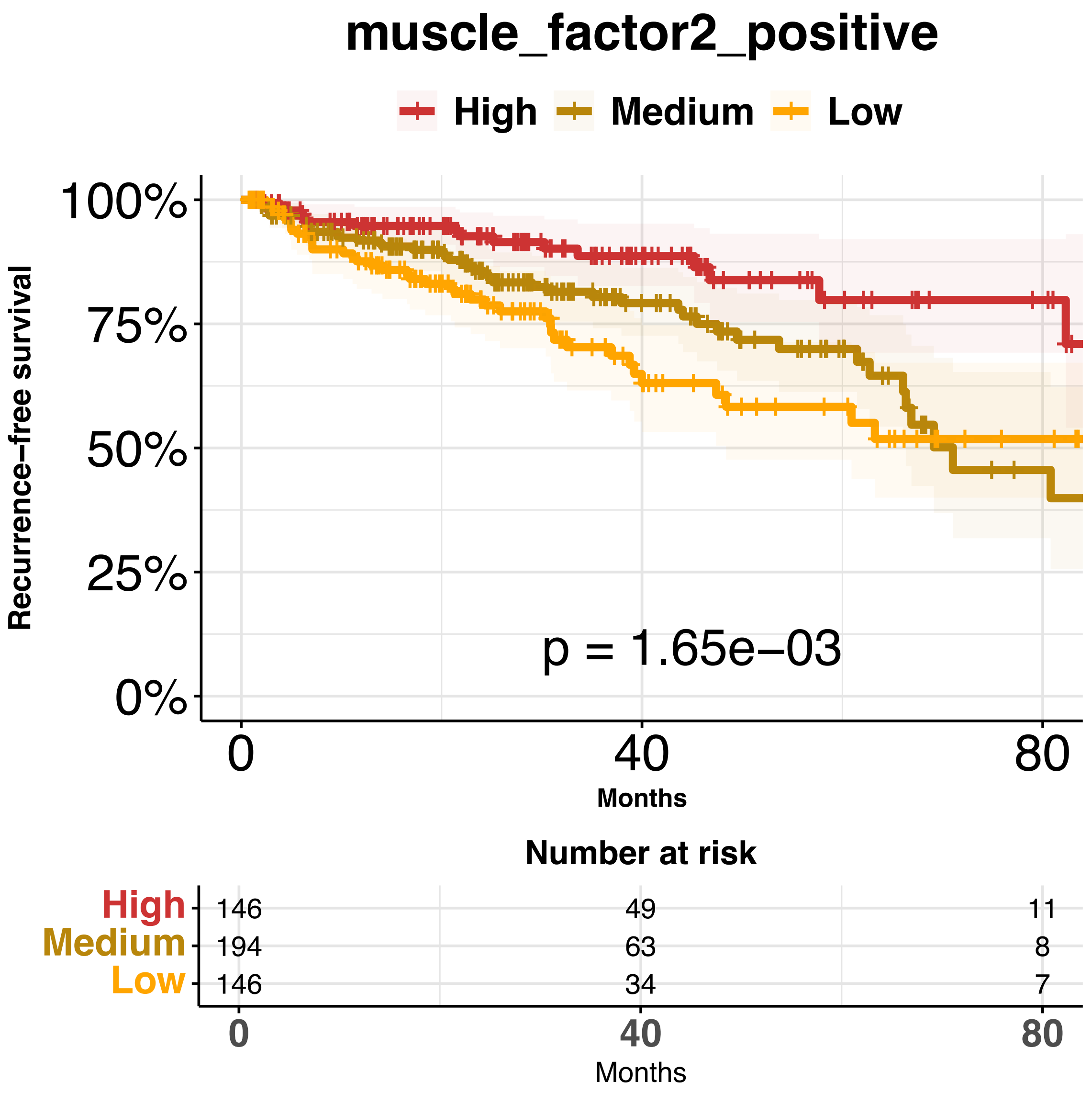


Supplementary Figure 9. Single-sample gene set enrichment for positive factor 2 weights for the biological theme “smooth muscle”.

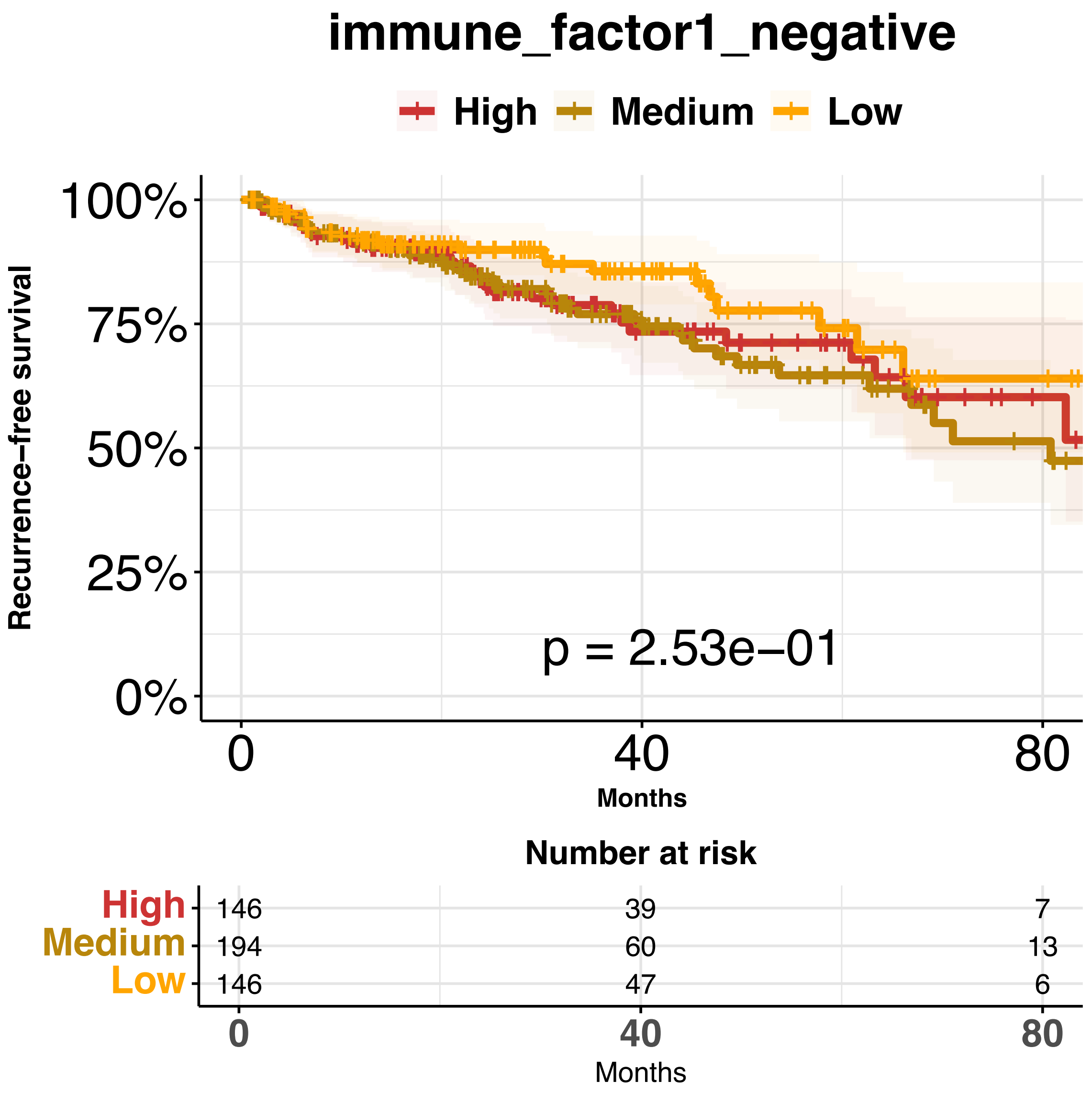


Supplementary Figure 10. Single-sample gene set enrichment for negative factor 1 weights for the biological theme “immune response”.

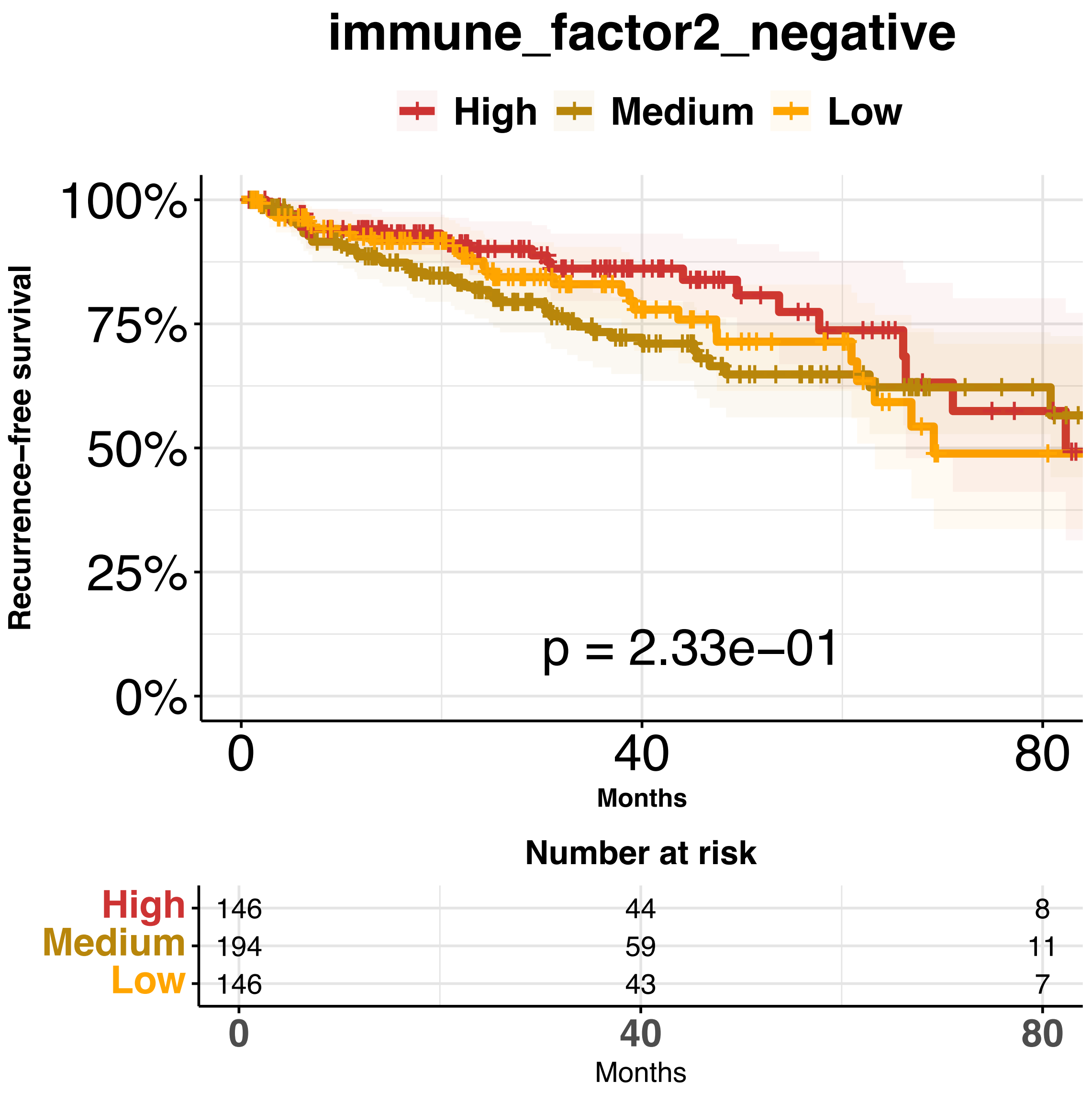


Supplementary Figure 11. Single-sample gene set enrichment for negative factor 2 weights for the biological theme “immune response”.

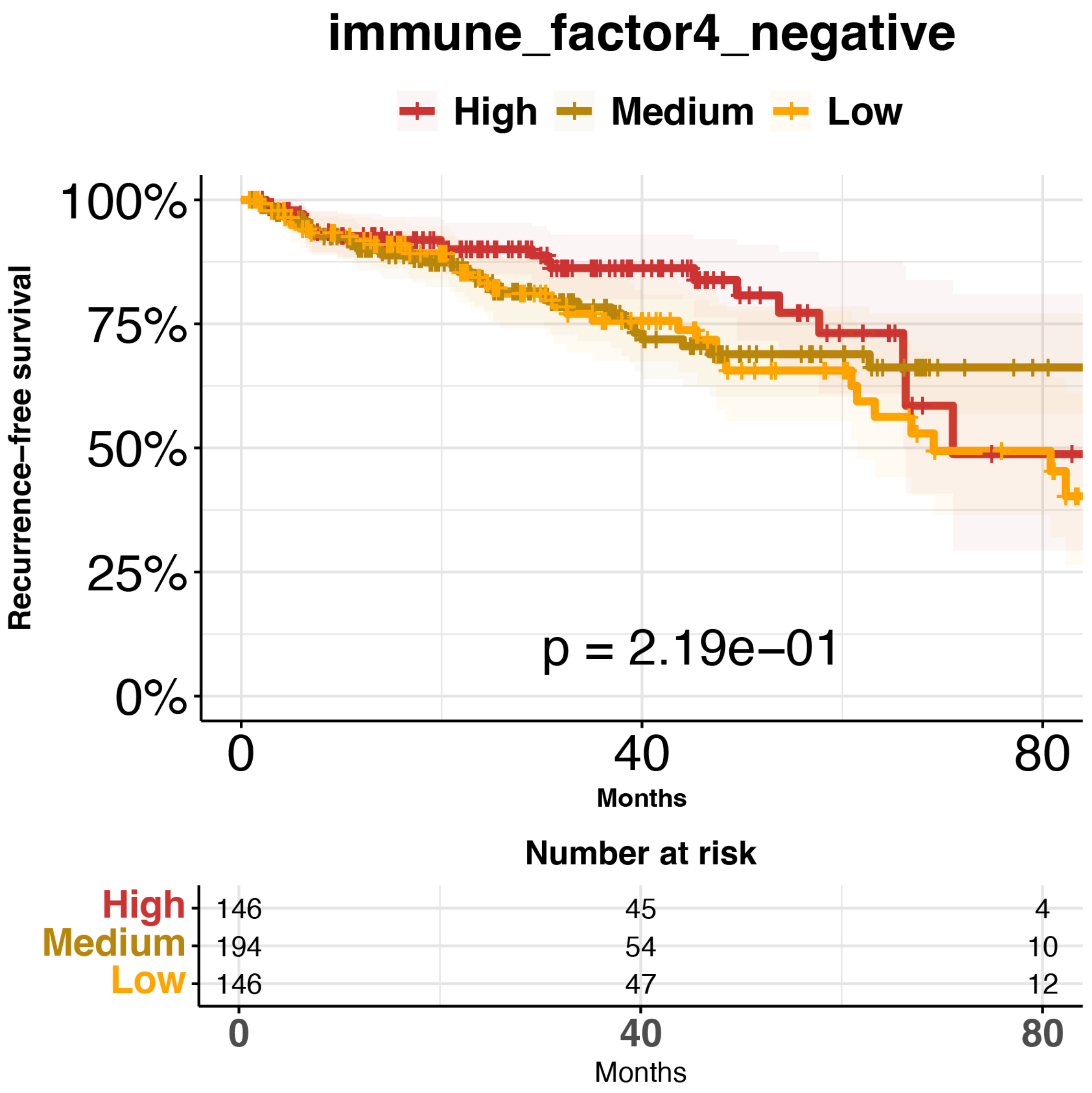


Supplementary Figure 12. Single-sample gene set enrichment for negative factor 4 weights for the biological theme "immune response".

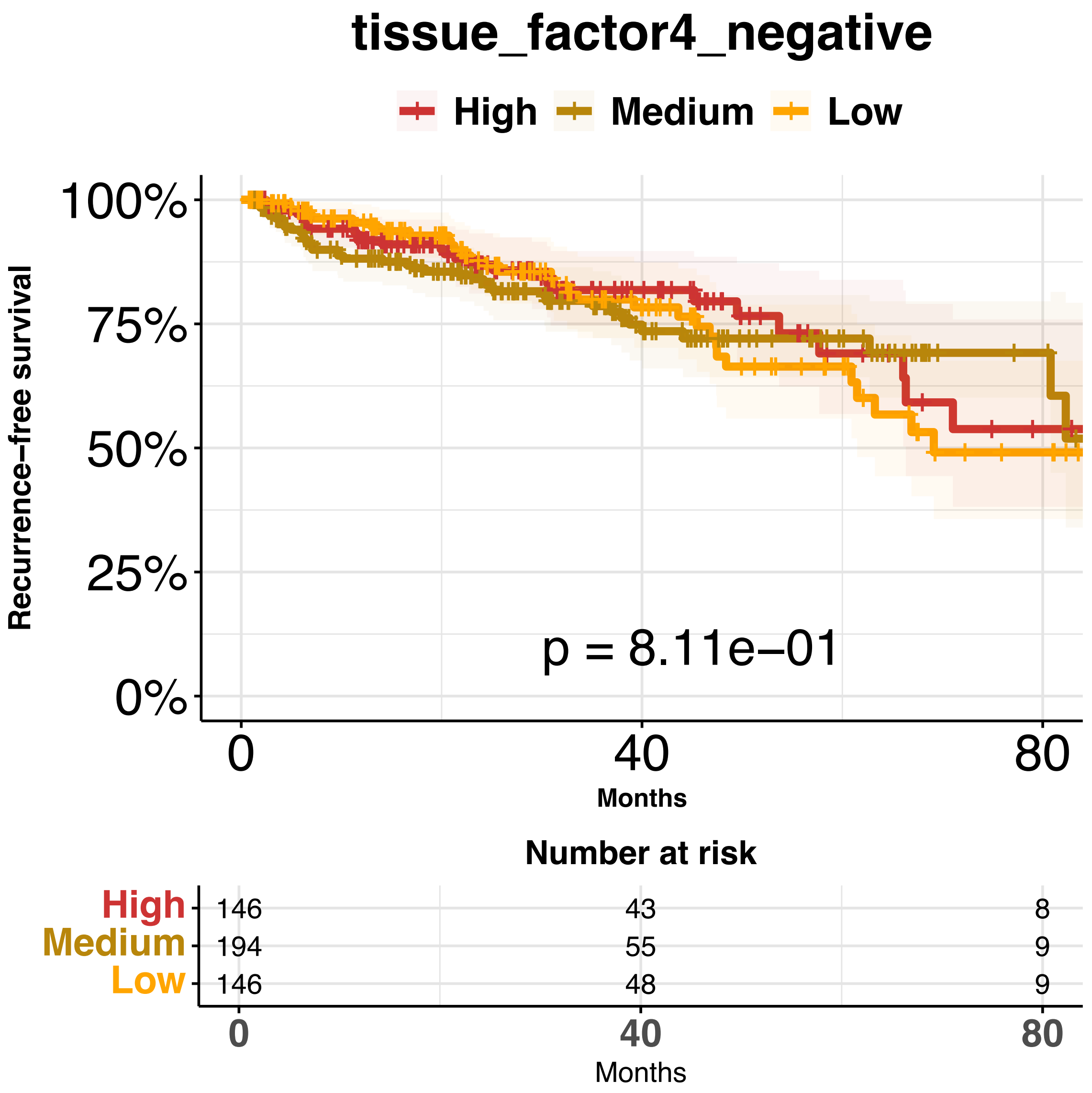


Supplementary Figure 13. Single-sample gene set enrichment for negative factor 4 weights for the biological theme "tissue morphology".

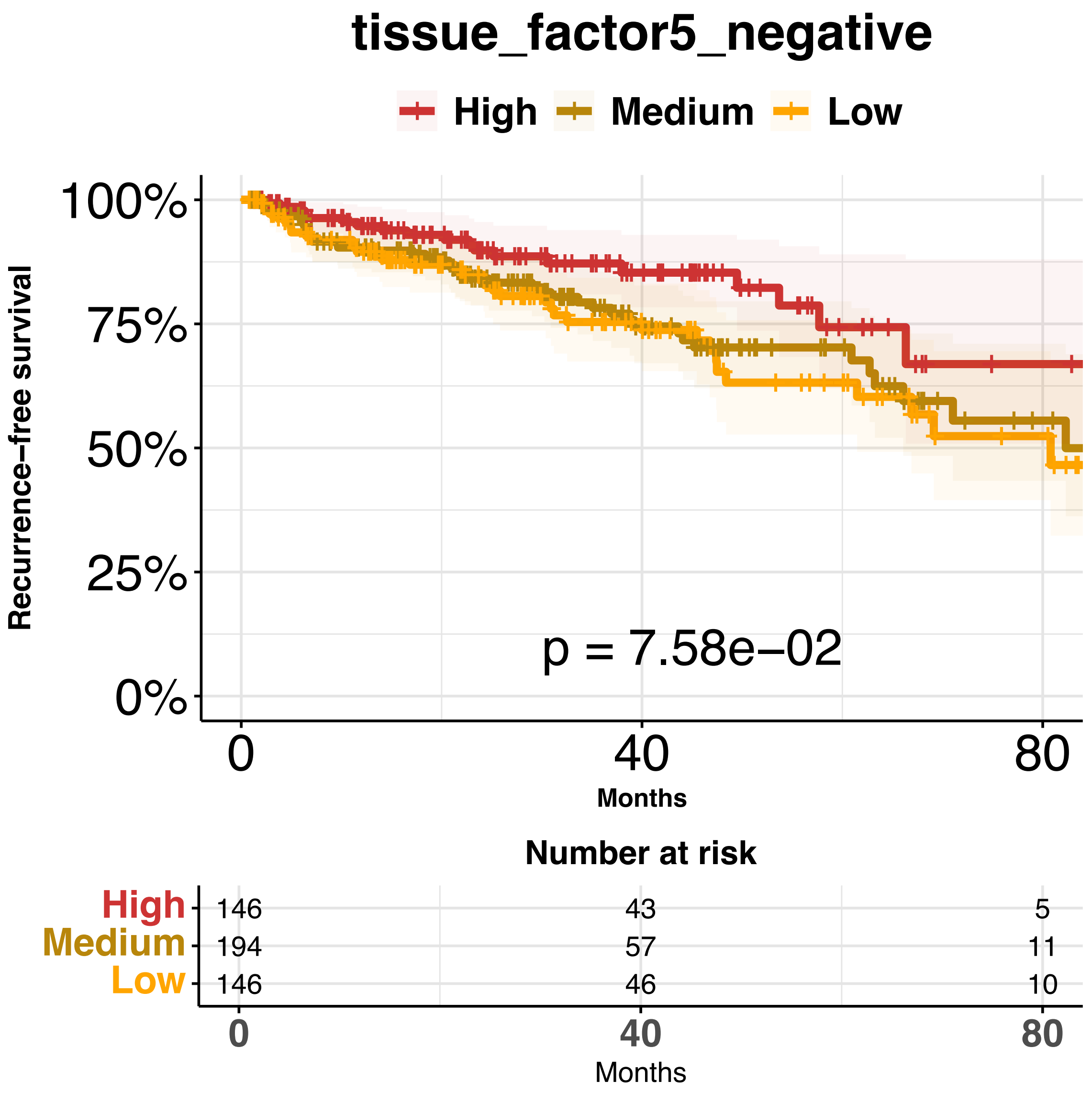


Supplementary Figure 14. Single-sample gene set enrichment for negative factor 5 weights for the biological theme “tissue morphology”.
